# Rapid and efficient oligodendrocyte differentiation from human pluripotent stem cells via dual inhibition of BMP and Notch signaling

**DOI:** 10.64898/2026.06.30.729930

**Authors:** Alessandro Evangelisti, Samantha M. Phillips, Johannes Jungverdorben, Ryan M Walsh, Youjun Wu, Vittoria Dickinson Bocchi, Ting Zhou, Lorenz Studer

**Affiliations:** Center for Stem Cell Biology, Memorial Sloan-Kettering Cancer Center, New York, NY 10065, USA; Developmental Biology Program, Memorial Sloan-Kettering Cancer Center, New York, NY 10065, USA; Neuroscience Program, Graduate School of Medical Sciences, Weill Cornell Medical College, New York, NY; Sloan Kettering Stem Cell Research Facility, Memorial Sloan-Kettering Cancer Center, New York, NY 10065

**Author notes:** Lead contact: Lorenz Studer.

## Abstract

The protracted timing required for oligodendrocyte differentiation from human pluripotent stem cells (hPSCs) has limited their use in disease modeling, drug screening, and cell therapy. In particular, the signals that drive oligodendrocyte specification and maturation after neural induction and ventral patterning remain poorly understood. Here, we present a protocol to derive human oligodendrocytes from hPSCs that is based solely on extrinsic cues, and we identify dual inhibition of BMP and Notch signaling as critical drivers of oligodendrocyte commitment and maturation. By day 42 of differentiation, up to 70% of the cells are positive for the oligodendrocyte marker O4, with minimal astrocyte contamination, and show robust expression of mature myelin markers including MBP, MOG, and MAG. These hPSC-derived oligodendrocytes closely match the molecular identity of primary fetal human oligodendrocytes as assessed by single-cell RNA sequencing and are functional as shown by *in vitro* myelination assays. In addition to the rapid generation of myelinating oligodendrocytes, the new protocol can be modularly adapted for the efficient production of PDGFRα+ oligodendrocyte precursors or mixed glial populations containing AQP4+ astrocytes, thereby providing a cellular toolbox for the study of human glial lineages in translational applications.

## Introduction

Oligodendrocytes represent one of the most abundant glial cell types^1^ and are responsible for CNS myelination. Myelination involves the formation of lipid-rich membrane structures that electrically insulate axons to enable saltatory conduction of action potentials critical for rapid neuronal communication^2,3^. Beyond myelination, oligodendrocytes play important roles in axonal metabolic support and ion homeostasis. In addition, oligodendrocyte progenitor cells (OPCs) are capable of direct synaptic interactions with neurons^4–7^. Therefore, the oligodendrocyte lineage embodies a dynamic and multifunctional cellular system essential for both neural circuit function and homeostasis. During forebrain development, an initial wave of OPCs arises from the medial ganglionic eminence (MGE) and the anterior entopeduncular area (AEP) around E12.5 in the mouse and around 8-10 weeks p.c. in human development^8–10^. In the developing human spinal cord, OPCs emerge from the ventral pMN domain of the neural tube around 7-9 weeks post conception (p.c.)^11,12^. The pMN domain represents a tightly restricted region defined by the co-expression of NKX6.1 and OLIG2 transcription factors^13–15^. This domain sequentially generates motor neurons followed by OPC lineage cells, which subsequently migrate away from the ventricular zone and differentiate and mature into myelinating oligodendrocytes^10,11,16–18^.

Numerous attempts to generate hPSC-derived oligodendrocytes *in vitro* have been made, building on a strong foundation of early embryonic development studies^19–23^. However, challenges with protocol consistency/reproducibility, protracted differentiation periods (often in excess of several months *in vitro*), low yields of mature cells, and the use of poorly defined media components have limited the use of oligodendrocytes across various translational applications *in vitro* and *in vivo*. Both committed O4+ oligodendrocytes and more mature MBP+ cells typically appear only at late stages of current *in vitro* differentiation protocols; this likely stems from the fact that oligodendrocytes are the final cell type to appear in their mature form in the CNS during *in vivo* development^24,25^, and thus there is a corresponding protracted differentiation time *in vitro*. Furthermore, while the induction of ventral precursor domains that give rise to oligodendrocyte lineage cells is well understood, the signals that further drive oligodendrocyte specification and maturation remain poorly understood. In addition, most current protocols lack unbiased validation of cell identity against scRNAseq profiles of primary oligodendrocytes and related lineages from the emerging sets of human fetal brain atlases.

Here, we present a novel, fast, and highly reproducible protocol to generate human oligodendrocytes from hPSCs that does not require ectopic expression of transcription factors such as SOX9 or SOX10^26,27^. By day 42 (6 weeks), we obtain up to 70% O4-positive cells, most of which are positive for myelin markers such as *MBP, MOG*, and *MAG*. The key innovation of this protocol is the combined inhibition of BMP and Notch signaling, which suppresses astrocyte differentiation and promotes accelerated oligodendrocyte commitment and maturation. Using scRNA sequencing combined with mechanistic studies, we describe the developmental trajectory from ventral precursor induction to OPC establishment and oligodendrocyte maturation, and we define the signaling pathways governing temporal cell fate transitions and identity acquisition. Leveraging these developmental insights, we have developed modular differentiation conditions to rapidly and efficiently derive oligodendrocytes and related lineages relevant to translational applications including: (1) an expandable PDGFRα+ OPC population, (2) mature oligodendrocytes expressing MBP, and (3) mixed glial cell populations comprising both astrocytes and oligodendrocytes.

## Results

### Overview of oligodendrocyte induction protocol - from pMN domain to mature oligodendrocytes

To track human OPC induction and oligodendrocyte differentiation, we first developed a dual-reporter hPSC line for OLIG2 and SOX10 expression (SOX10::H2B-GFP; OLIG2::H2B-tdTom) as detailed (**Fig. S1** and ^28^). We established a 3D organoid-based differentiation protocol to apply an optimized sequence of extrinsic patterning cues (**Fig. 1A,B**) under conditions that promote cell-to-cell interactions during differentiation^5,29^. Within 42 days of differentiation, cells progress from hPSCs to the pMN lineage, glia-competent precursors, oligodendrocyte-biased lineages and OPCs, and ultimately to mature, myelinating oligodendrocytes marked by myelin basic protein (MBP) (**Fig. 1B, C, S2A, B**). During the first part of the protocol (day 0-26 of differentiation), cells reach a glial competent state that represents a bi-potent precursor characterized by co-expression of *SOX9* and *NFIA*. To optimize expression of OLIG1 and OLIG2 transcription factors (TFs), crucial for oligodendrocyte lineage development within the glial-competent lineage, we implemented three key modifications to established protocols. As early as day 6, we introduced exposure to **(1) RA agonists** that target both RARα and RXR receptors with high affinity. The use of RA agonists instead of all-trans retinoic acid improved the induction of the pMN domain while minimizing the presence of dorsal and ventral off-target populations, including contaminating FOXA2+ floorplate-like cells. At day 12 of differentiation, we added **(2) FGF18** to inhibit neurogenesis from pMN precursors which, prior to the induction of gliogenesis, produce spinal motor neurons during normal *in vivo* development. Furthermore, FGF18 promoted the proliferation of glial-competent cells that retain expression of OLIG2. At day 22, we included a potent sonic hedgehog inhibitor, **(3) cyclopamine**, to suppress any remaining FOXA2+ floorplate-like progenitors.

**Figure 1.**
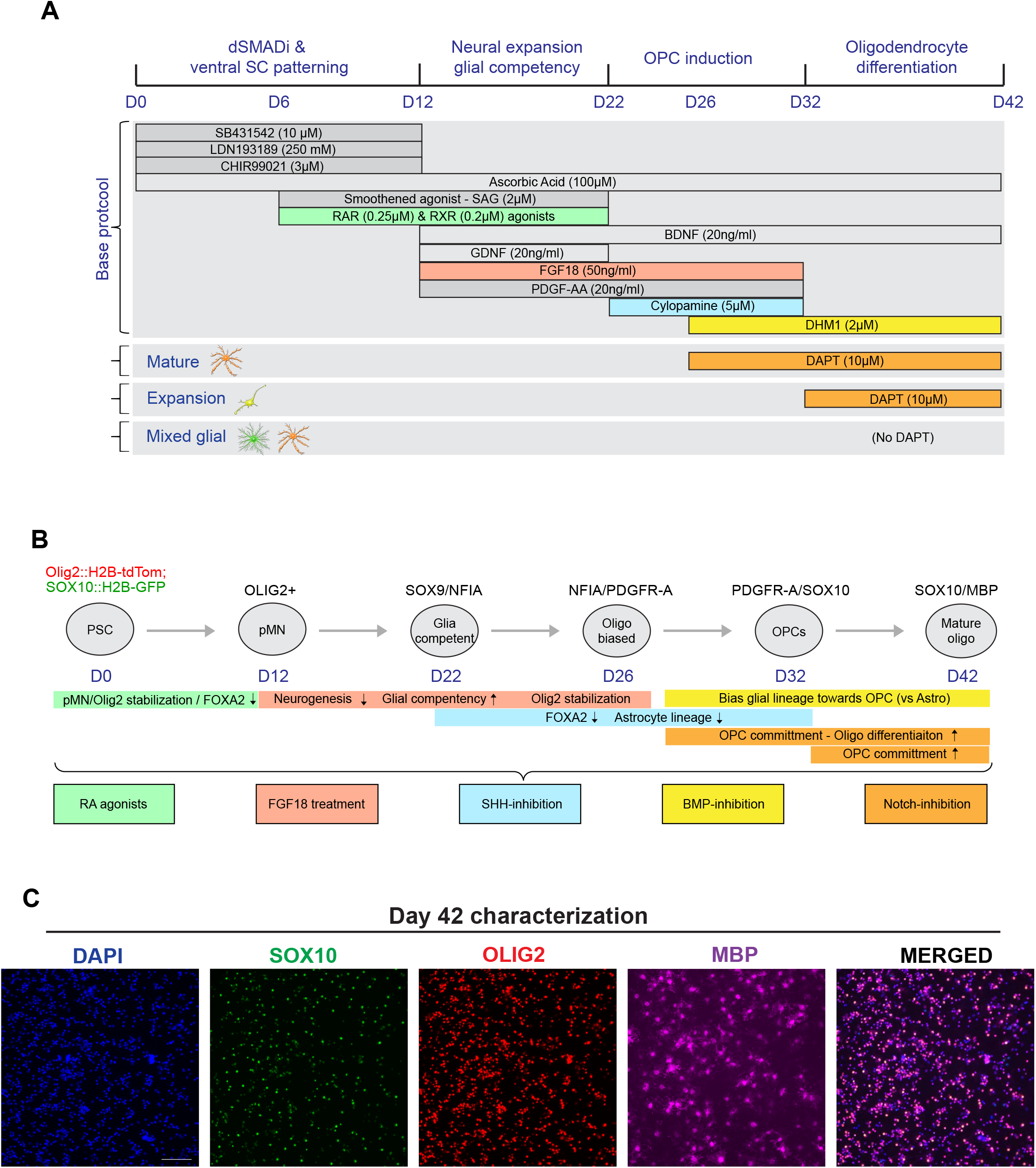
Overview of oligodendrocyte induction protocol - from pMN domain to mature oligodendrocytes. (A) Schematic illustration of the 42-day organoid-based protocol that enables rapid differentiation of human PSCs to mixed-glial, OPC, and mature oligodendrocyte populations. B) Developmental model showing the key modifications in modulating signaling pathways for oligodendrocyte specification and differentiation - from spinal cord ventral pMN domain induction to generating mature oligodendroglia along the 42-day protocol. (C) Representative immunofluorescence images show cells positive for OLIG2, SOX10, and myelin basic protein (MBP) at day 42. Scale bar: 100μm.

During the second phase of the protocol (day 26-42 of differentiation), we established differentiation conditions involving dual inhibition of BMP and NOTCHi to derive either proliferating OPCs, mature oligodendrocytes or mixed glial lineages comprising both mature oligodendrocytes and astrocytes (**Fig. 1A**). For all three conditions, day 26 precursors were treated with **(4) DMH1**, a strong BMP signaling inhibitor to prevent the shift from bi-potent to astrocytic committed fate. The key driver for further oligodendrocyte lineage specification is the inhibition of Notch signaling via treatment with the γ-secretase inhibitor **(5) DAPT**. Sustained NOTCHi (day 26-42) promoted exit from bi-potent towards SOX10+ committed oligodendrocyte lineage and further differentiation into mature MBP+ cells. In contrast, a shorter or delayed pulse of NOTCHi promoted the emergence of PDGFRα+ OPCs (**Fig. 1B**). Finally, the omission of NOTCHi from day 26-42 in the presence of DMH1 promoted differentiation into a mixed population of AQP4+ astrocytes and SOX10+ oligodendrocytes. Importantly, our mature oligodendrocyte protocol, based on dual inhibition of BMP and Notch, enabled a highly efficient output of mature oligodendrocytes in the absence of astrocytes within 6 weeks of hPSC differentiation (**Fig. 1C**).

### From pMN domain induction to bipotent, glial-competent precursors – RA agonist, sonic hedgehog inhibitor and FGF18 treatment reduce FOXA2+ astrocytic lineage and prevent OLIG2 loss

The pMN domain is highly restricted within the ventral developing spinal cord, and pMN induction requires the correct balance of dorsoventral (D/V) patterning signaling. Retinoic acid acts within the ventral spinal cord as a putative dorsalizing signal^30^ to balance the extent of sonic hedgehog (SHH)-mediated ventralization and the combined treatment with retinoic acid and SHH can “lock” cells within the pMN domain^31,32^. In our current protocol, we confirm by single cell RNA sequencing that by day 12, our cells express the major transcription factors^14,33^ that define the pMN domain (**Fig. 2A, left panel**). This includes expression of NKX2.2, a classic marker of the more ventral p3 domain in the mouse but shown to give rise to a subset of late born motor neurons during human development^34^. While NKX2.2 is typically restricted to the p3 domain during neurogenesis, there is dorsal expansion of NKX2.2 during gliogenesis into the pMN domain, and pMN domain progenitors that go on to make oligodendrocytes express both Olig2 and NKX2.2^14,35,36^.

**Figure 2.**
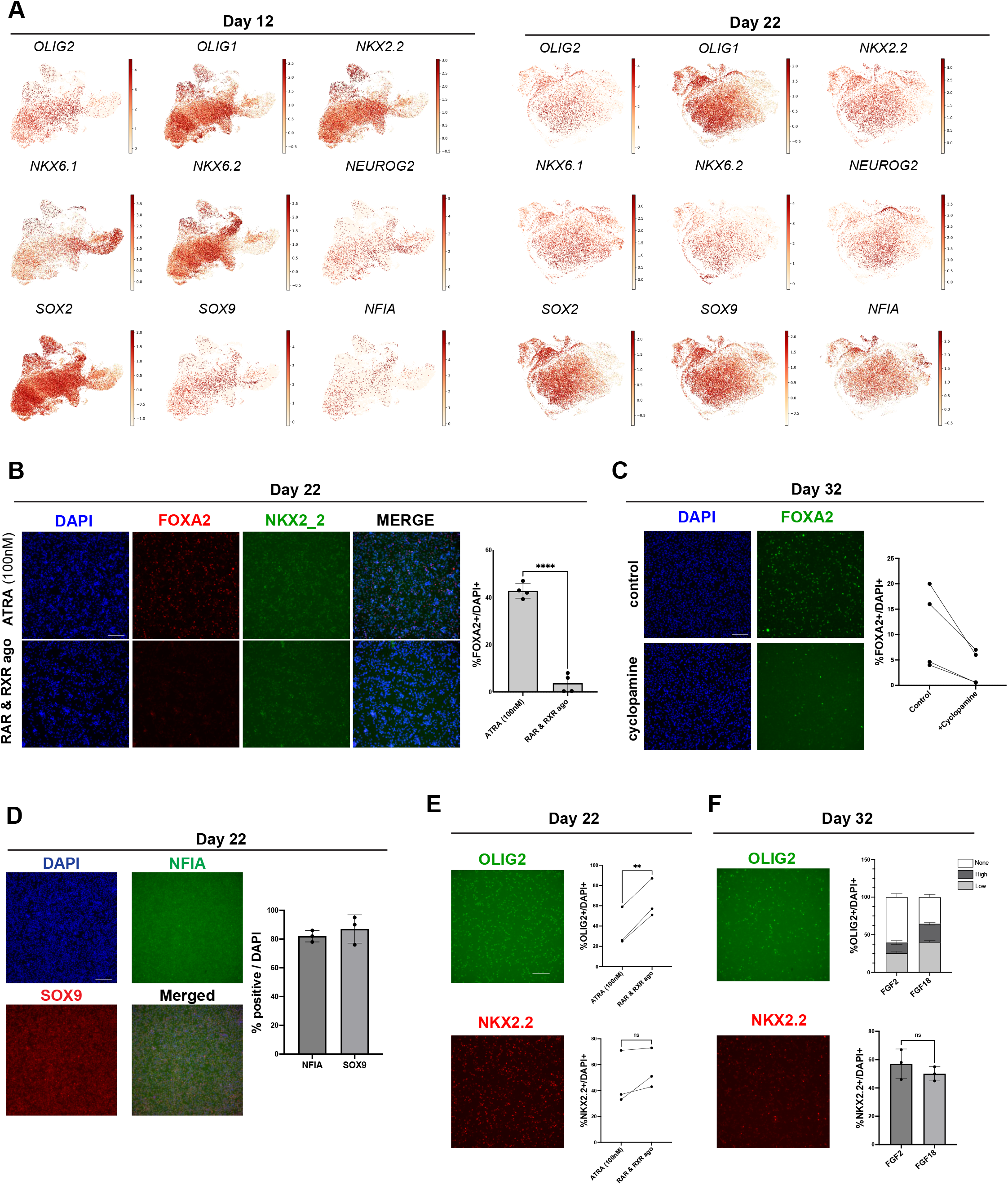
From pMN domain induction to bipotent, glial-competent precursors –RA agonist, sonic hedgehog inhibitor and FGF18 treatment to reduce FOXA2+ astrocytic lineage and prevent OLIG2 loss. (A) Single gene expression UMAPs from scRNAseq data show the correct patterning of the pMN domain at day 12 based on the expression of major transcription factors such as *OLIG2* and *NKX6*-*2*, and the rapid conversion to a glial competent state, *NFIA-SOX9*, by day 22. (B) Significant downregulation of FOXA2+ floor plate-like cells by day 22 using retinoic acid agonists (RAR and RXR) instead of the more commonly used all-trans retinoic acid (ATRA) treatment. (C) Further inhibition of FOXA2+ floor plate cells by blocking endogenous sonic hedgehog signaling with *cyclopamine* from day 22 to day 32. D) Representative immunofluorescence images of the glia competent state show the expression of *NFIA*-*SOX9* achieving induction of these markers in >80% of the culture by day 22. (E) OLIG2 expression significantly increased in RA agonist-treated cultures at day 22 compared to ATRA treatment, while *NKX2-2* did not significantly change. (F) OLIG2 downregulation by day 32 was partly rescued by treating the cells from day 22 to day 32 with FGF18 instead of more broadly used FGF2, while *NKX2-2* did not significantly change under these conditions. OLIG2 expression was quantified based on high and low expression profiles. Scale bar: 100μm

By day 22, most of the cells co-express *NFIA* and *SOX9*, while retaining *OLIG1/OLIG2* expression suggesting that they adopt glial competency and bi-potential ability towards either astrocytes or oligodendrocytes (**Fig. 2A, right panel**). In published protocols, RA signaling is typically activated by exposure to ATRA (all-trans retinoic acid). However, when comparing ATRA versus RA agonist treatment, we observed the emergence of a FOXA2+ off-target population, co-expressing NKX2.2 (**Fig. 2B**) in ATRA treated cultures. We hypothesized that these FOXA2+/NKX2.2+ precursors may bias lineage specification towards a later-stage astrocytic lineage. Astrocytes emerge as a common “off-target” of all major extrinsic factor-based oligodendrocyte differentiation protocols^37,38^, and minimizing FoxA2 contaminants may improve oligodendrocyte yield at the expense of the production of off-target astrocytes. In addition to possibly becoming astrocytes, it is also possible that these FOXA2+/NKX2.2+ precursors would become interneurons, as cells with that signature are also expressed in the ventral-most cells of the p3 domain in mice, which can become Sim1+ V3 interneurons^39^. We found that RA agonists that target both the RAR and RXR receptors significantly reduce differentiation into FOXA2+ cells by day 22 as compared to ATRA treatment (**Fig. 2B, Fig. S3A**). Nevertheless, even in RA agonist-treated cultures, we noticed proliferation of a small population of FOXA2+ precursors from day 22 – 32. We hypothesized that endogenous sonic hedgehog signaling might contribute to continued proliferation of FOXA2+ contaminants. Accordingly, we tested the impact of exposing RA agonist-treated cells to cyclopamine, a potent SHH inhibitor, from day 22 – 32. Cyclopamine treatment eliminated nearly all FOXA2+ contaminants (**Fig. 2C**) by day 32 of differentiation while also limiting ISL1+ motor neuron development (**Fig. S3B**). However, despite eliminating most FOXA2+ cells and inducing a significant increase in OLIG2+ cells at day 22 in organoids exposed to RA agonists (**Fig. 2E**), we noticed a downregulation of OLIG2 expression from day 22 to day 32 (**Fig. 2E, F**). Instead of the widely used FGF2 treatment to promote OLIG2 expression, we treated the cells with FGF18 (from day 12 – 32). FGF18 treatment performed better than FGF2 in stabilizing pMN-derived lineages and OLIG2 expression (**Fig. 2F, S3C**) and enabled the efficient expansion of a bi-potent oligodendrocyte-biased precursor lineage. Additionally, cells remain strongly positive for *OLIG1* and progressively transition to PDGFRα+ progenitors and committed SOX10+ oligodendrocytes (**Fig. S3D, E**).

### BMP inhibition prevents astrocyte commitment and promotes glia-competent state capable of oligodendrocyte lineage specification

Glial competent, bi-potent cells at day 26 are characterized by the expression of the transcription factors *SOX9* and *NFIA* in combination with OLIG1/OLIG2. We reasoned that biasing cells toward an oligodendrocyte fate will first require maintaining high levels of OLIG1/OLIG2 expression (**Fig. 2F, S3D**). Second, cells that retain high levels of OLIG1/OLIG2 expression should progress in a time-dependent manner towards oligodendrocyte lineage commitment characterized by the expression of SOX10. We found that strong inhibition of BMP signaling directed bi-potent precursors towards NFIA/SOX2/PDGFRα oligodendrocyte lineage-biased precursors (**Fig. 3A**), as further illustrated by the emergence of SOX10/PDGFRα/OLIG2 population at day 42 of the differentiation (**Fig. 3B**). A potential mechanism by which BMPi may bias towards oligodendrocyte lineage has been described in past studies focusing on astrocyte development. In fact, previous work in primary neural precursors reported that activating the BMP pathway induces differentiation into astrocytes both *in vivo* and *in vitro*^40–42^. However, these studies, while showing a decrease in oligodendrocytes when cells are treated with BMPs, did not directly address whether there is a concomitant increase in oligodendrocyte lineage specification upon inhibiting BMP signaling.

**Figure 3.**
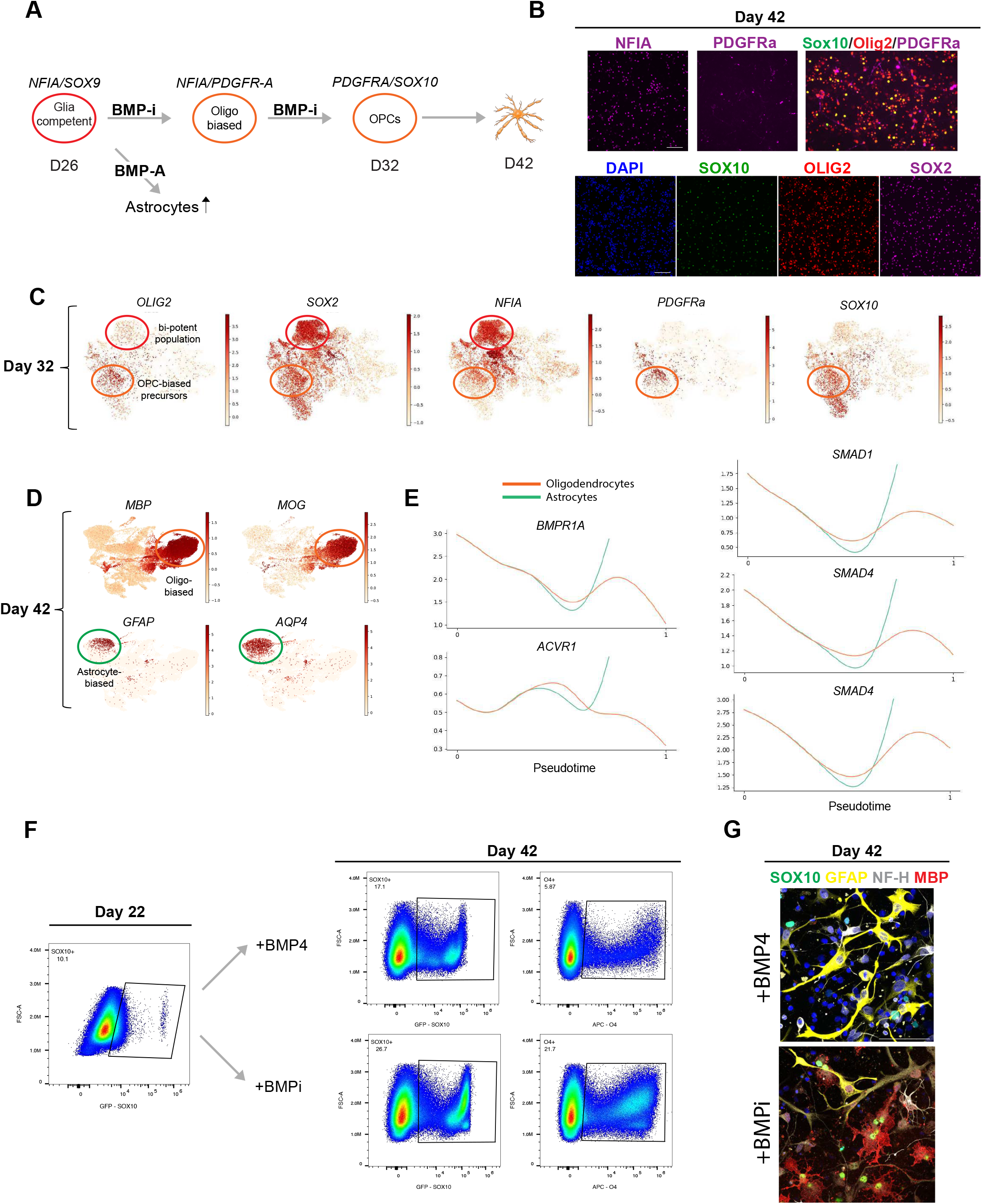
BMP inhibition prevents astrocyte commitment and promotes glia-competent state capable of oligodendrocyte lineage specification. (A) Schematic illustration depicting the proposed role of BMP inhibition in biasing a glia bi-potent population towards oligodendrocyte progenitor cells and against astrocyte differentiation. (B) Representative immunofluorescence images show a strong induction of a glial-biased *OLIG2-SOX2-NFIA* bipotent population, which gives rise to OPC-biased progenitors positive for PDGFRα and low in *SOX10*. (C) ScRNAseq data from day 32 defines two distinct populations: a bi-potent glial biased at first, which switches to OPC precursors fate, positive for OLIG2-*PDGFRα-low SOX10*. (D) BMPi alone allows the differentiation to both mature *MOG/MBP* oligodendrocytes as well as to GFAP-AQP4 astrocytes (*mixed glial* condition). (E) Cell trajectory analysis using Palantir shows the upregulation of BMP signaling (both receptors and modulators) in astrocytes and concomitant downregulation in oligodendrocyte development. (F) Flow based analysis of BMP4 versus DMH1 (BMPi) treated cultures. BMP4 treatment from day 26 to day 42 resulted in a decrease in SOX10 by day 32, and a significant drop in O4 population by day 42 compared to DMH1 treated cells. Scale bar: 100μm. G) Immunocytochemistry confirms induction of GFAP protein and lack of MBP expression at day 42 following BMP4 treatment.

By scRNAseq at day 32, using our *mixed-glial* protocol involving BMPi only, we show the presence of two distinct populations. There is a putative bi-potent *SOX2-NFIA-OLIG2*-*positive population*, and a population biased towards OPC lineage expressing *NFIA-OLIG2-PDGFRα* and initiating expression of *SOX10* (**Fig. 3C**). By day 43, these OPCs presumably differentiated into mature oligodendrocytes characterized by major myelin components, *MBP* and *MOG*, while the previously unbiased progenitors transitioned into differentiated astrocytes positive for *GFAP* and *AQP4* (**Fig. 3D**). We further tracked key components of the BMP signaling pathway in our scRNAseq dataset throughout all differentiation time points (days 12, 22, 32, and 43). We applied the lineage trajectory tool *Palantir*^43^ to track the predicted developmental trajectory of cells identified at endpoint as astrocytes or oligodendrocytes (day 43). We observed a distinct trajectory along pseudotime with early, marked upregulation of BMP signaling molecules in the astrocyte lineage (green line). In contrast, the oligodendrocyte-lineage trajectory (orange line) showed a marked time-dependent decrease in BMP signaling pathway components (**Fig. 3E**). We next tested whether we could manipulate the differentiation propensity of early bi-potent precursors towards astrocytic lineages. Treatment with BMP4, from day 22 onwards, decreased the emergence of SOX10+ cells and reduced the percentage of O4+ cells by day 42 by about 4-fold (**Fig. 3F**). Conversely, BMP4 treatment induced GFAP+ astrocytes and largely blocked the differentiation into MBP+ oligodendrocytes (**Fig. 3G**).

### Notch inhibition triggers oligodendrocyte specification in bi-potent precursors and promotes transition from OPCs into mature oligodendrocytes

While BMPi reduced the propensity for astrocyte differentiation and enabled the time-dependent transition into oligodendrocyte lineage cells, it did not directly trigger oligodendrocyte induction. We hypothesized that time-dependent oligodendrocyte differentiation may involve a feedback mechanism within pools of bipotent progenitor cells that controls the progressive differentiation. During neuronal differentiation, inhibition of Notch signaling among neuronal precursors can trigger neuronal differentiation^44–46^. To assess whether oligodendrocyte differentiation may be dependent on a similar mechanism, we treated bipotent precursor cells at day 26 with the γ-secretase inhibitor DAPT, a widely used Notch inhibitor. Flow cytometry analysis at day 42 demonstrated that DAPT treatment results in an increase in the percentage of oligodendrocyte lineage cells from 34.3% SOX10+, 22.4% O4+ and 13.0% O1+ for BMPi only (*mixed glial condition*), to 72.1% SOX10+, 64.3% O4+, 50.0% O1+ cells upon dual NOTCH- and BMPi (*mature condition*) (**Fig. 4A, S4A**). Interestingly, a delay in NOTCHi treatment by 6 days further increased the SOX10+ population (78.3% SOX10+ cells; **Fig. S4B**) but led to a reduced percentage of O4+ oligodendrocytes (*expansion condition*). We further noticed that the delay of NOTCHi resulted in a population of cycling OPCs expressing PDGFRα at day 42 (**Fig. S4C**). To assess whether we can further expand PDGFRα OPCs, we treated our organoids with early NOTCHi from day 26-32 followed by DAPT withdrawal in the presence of FGF18 and DMH1. In addition, we added recombinant PDGF-AA, produced in CHO cells. CHO-PDGF-AA is more potent than E. coli expressed PDGF-AA. Under these conditions we saw a robust increase in the percentage of PDGFRα expressing cells by day 42 (**Fig. S4C**). In contrast, transition to mature oligodendrocytes upon DAPT treatment was further confirmed by increased percentage of MBP+ and PLP1+ cells in the *mature* protocol compared to conditions without DAPT (**Fig. S4D**) We performed scRNAseq analysis at day 43 for each of these three conditions (**Fig. 4B**) including no NOTCHi (*mixed glial*), late NOTCHi (*expansion*), continued NOTCHi (*mature*). The distinct transcriptional identities of the three populations were marked by *AQP4, PDGFRα*, and *MBP* expression respectively (**Fig. 4C**). Trajectory analysis along pseudotime using *Palantir*^43^ showed that developmental progression toward either astrocyte or oligodendrocyte terminal states followed symmetric but opposing patterns of Notch regulation. Astrocytic fate was linked to a strong, time-dependent increase in the expression of Notch signaling targets (*HES1* and *HES5*) while the oligodendrocyte lineage trajectory was characterized by a delayed decrease in the expression of *HES1* and *HES5* (**Fig. 4D**).

**Figure 4.**
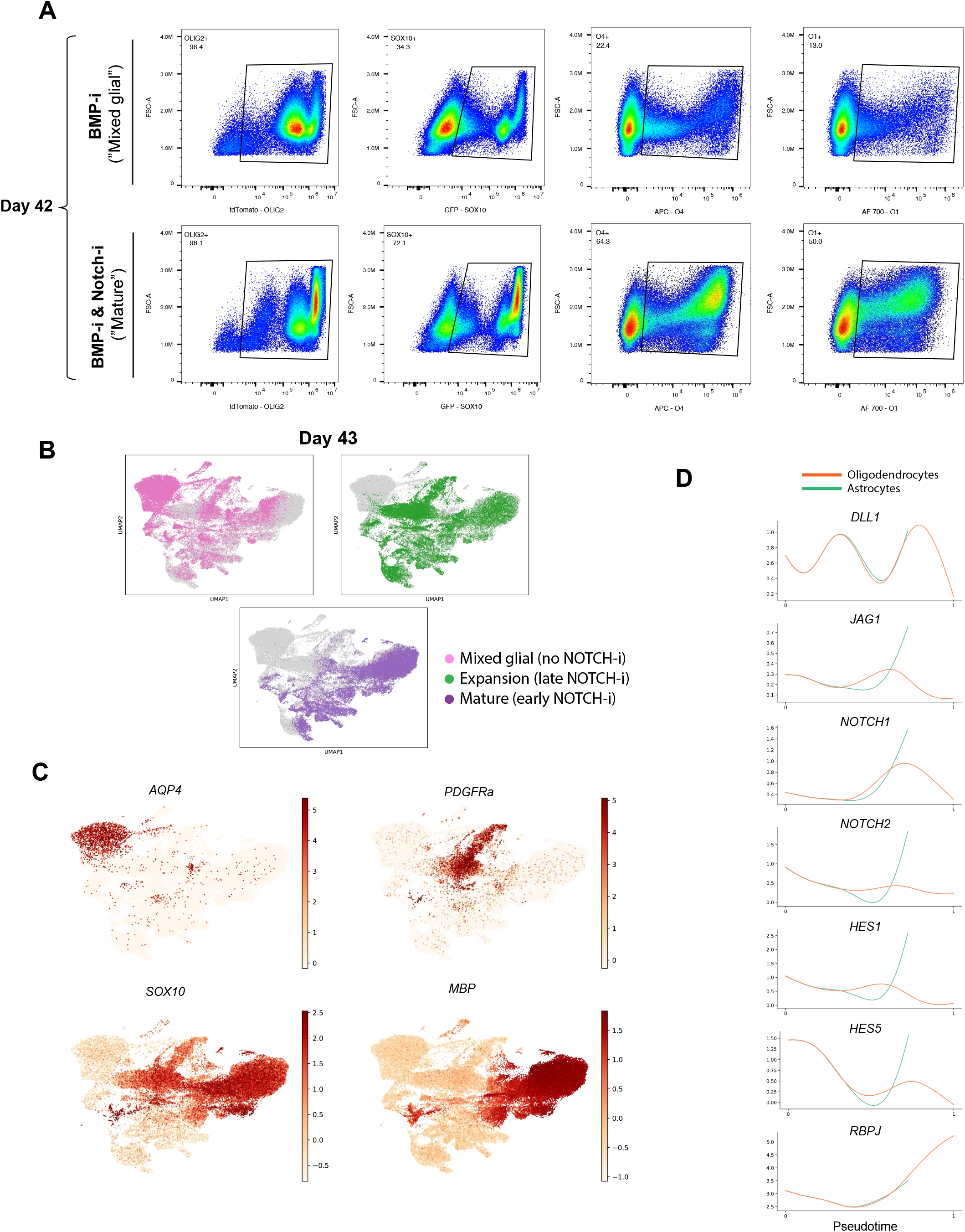
Notch inhibition triggers oligodendrocyte specification in bi-potent precursors and promotes transition from OPCs into mature oligodendrocytes. (A) The inhibition of NOTCH together with BMPi at day 26 induces the transition of OPC-biased cells to an increased percentage of cells committed to SOX10 and O4-O1 pre-myelinating cell population compared to DMH1-only treated cells, as shown by flow cytometry analysis (22.4% O4 and 13.0% O1 in mixed glial, and 64.3% O4 and 50.0% O1 in mature). (B) Distribution of cells differentiated through the mixed-glial, expansion, and mature protocols (no NOTCHi, late-NOTCHi, or early NOTCHi, respectively) shown by scRNAseq data. (C) Single gene expression UMAPs define distinct cell identities as a function of NOTCH inhibition across the three protocols: the mixed-glial population mostly expresses astrocyte markers (AQP4), the expansion condition showed a strong upregulation of oligo progenitor cell markers (high PDGFRα), and the mature cluster showed an upregulation of mature oligodendrocyte markers such as MBP. (D) Trajectory analysis confirms the upregulation of the Notch signaling pathway in astrocytes and a strong downregulation in oligodendrocytes.

### Acquisition of astrocyte versus oligodendrocyte fate depends on BMP and Notch signaling

Our data suggest that the fate decision towards either astrocyte or oligodendrocyte terminal states is controlled by early BMP signaling, with BMPi required to limit astrocytic differentiation, followed by modulation of Notch signaling with NOTCHi inducing an OPC-biased lineage and promoting subsequent oligodendrocyte differentiation. To further support this model, we performed a more in-depth analysis of our scRNAseq data based on the *mixed-glial* differentiation paradigm that comprises a balanced proportion of both astrocytes and oligodendrocytes. We reconstructed the developmental pseudotime towards both terminal states using *Palantir*^43^. Our results indicate progressive time points of astrocyte and oligodendrocyte commitment (**Fig. 5A, S5A**). Terminal states were characterized by the induction of key astrocyte lineage markers including *CD44, GFAP*, and *AQP4* versus the induction of mature oligodendrocyte markers such as *MBP, MOG, and MAG* (**Fig. 5B, S5B**). Analysis of the glial competency factor NFIA suggested that induction is critical for both lineages. Around pseudo-time 0.5, all cells regardless of final identity, expressed high levels of *NFIA* (**Fig. 5C**). However, the oligodendrocyte lineage is characterized by a drastic decrease in *NFIA* expression following the initial induction, while astrocytic lineage retains high levels of expression towards the terminal state. *SOX9* follows a similar pattern of transient induction in oligodendrocyte lineage versus sustained expression in astrocyte lineages. Finally, strong induction of *OLIG2* and *PDGFRα* characterize the oligodendrocyte lineage with low and transient only expression within the astrocytic lineage (**Fig. 5D**). The induction of *OLIG2, PDGFRα* and other oligodendrocytes related genes, is delayed in the expansion protocol confirming the critical role of Notch downregulation in oligo lineage specification (**Fig. S5C, 5D**). Next, we focused our analysis on the bifurcation time point of glial subtype specification rather than terminal lineage reconstruction. We used *Genes2Genes*^47^ to compare gene level trajectory alignment between alternative fates^47^. We built an alignment cost landscape (where cost reflects how different gene expression trajectories are across cells), capturing the relationship between cell type composition and gene expression profiles. (**Fig. 5E**). Each bin (10 in total) corresponds to the diversity of transcriptional programs we can identify within our analysis. The heatmap indicates that from bin 5 and 6 of the oligodendrocytes, the “cost” or transcriptional program differences versus astrocytes greatly increased until cells fully committed to each fate, presumably around bin 10, where the cost reached the highest value, translated in the greatest transcriptional differences between the two lineages (**Fig. 5E**). By mapping the temporal information of the corresponding scRNAseq datasets, we observed that bins 5 and 6 correspond largely to day 22 of differentiation (see color-coded legend). This data suggests that glia-competent cells at this early differentiation time point already exhibit an intrinsic bias towards either cell fate (**Fig. 5E**). Single gene expression plots along pseudo-time showed the induction of BMP and Notch signaling (*SMAD1-4* and *HES1-5*, respectively) at bin 5 and correlated with the induction *NFIA* and *SOX9* within the astrocyte lineage, while the oligodendrocyte trajectory showed only limited and delayed induction of these pathways (**Fig. 5F**). These data further support a paradigm involving dual inhibition of BMP and Notch signaling pathways as a critical feature in driving oligodendrocyte lineage bias and differentiation, while reducing astrocytic commitment.

**Figure 5.**
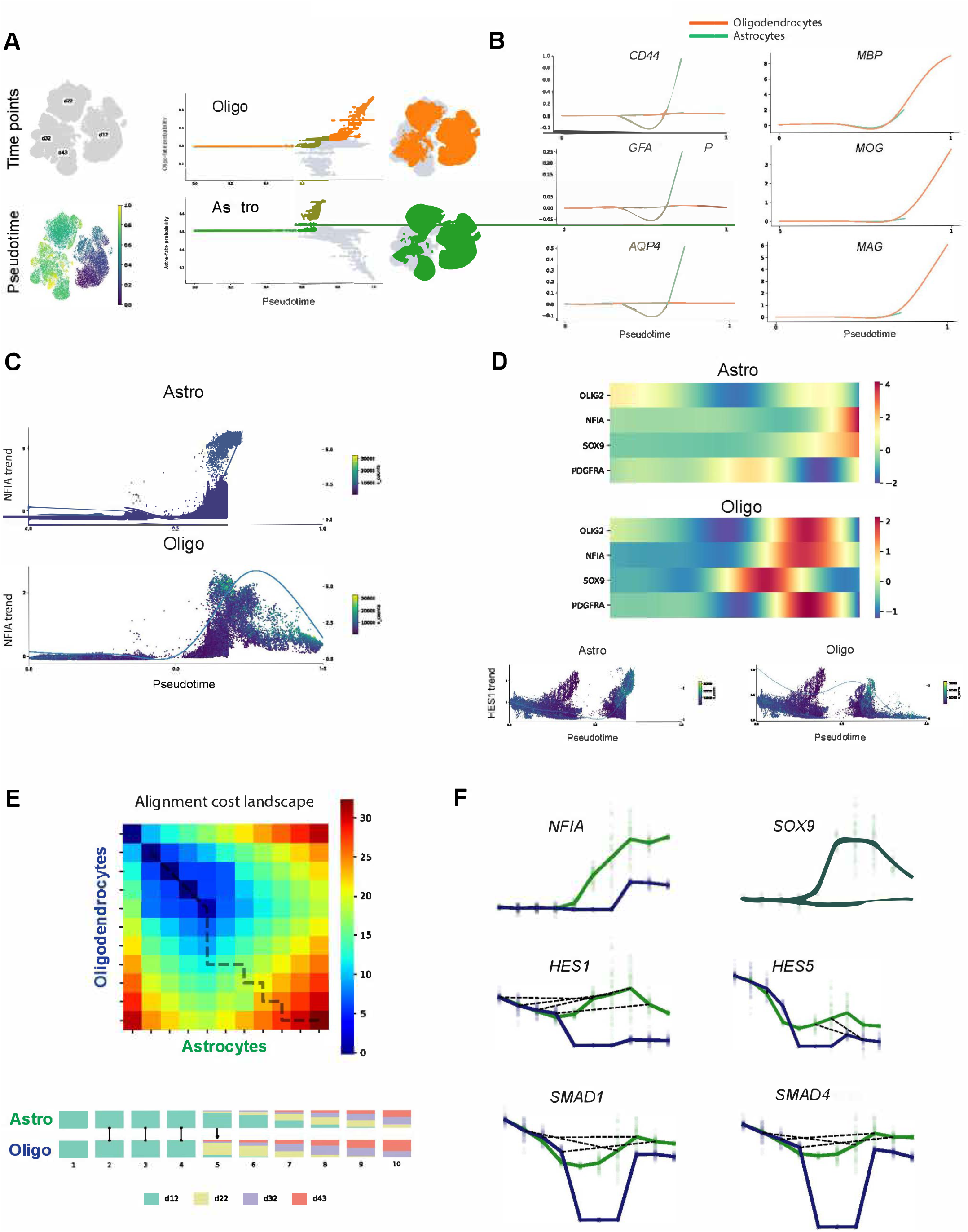
Acquisition of astrocyte versus oligodendrocyte fate depends on BMP and Notch signaling. (A) Pseudotime trajectory illustrating oligodendrocyte and astrocyte lineage progression (B) Gene trends graphs define the two distinct glial populations derived from the mixed-glial protocol. (C) *NFIA* expression peaks during early glia development, decreases drastically as cells become committed OPCs, while steeply increases as cells differentiate to mature *AQP4* astrocytes. (D) Gene expression heatmaps describe the role of major transcription factors involved in the switch between oligodendrocyte and astrocyte cell fates. (Bottom) *HES1* gene trend shows a strong correlation of astrocyte versus oligodendrocyte commitment by NOTCH activation or inhibition, respectively. (E) Heatmap illustrates gene level trajectory alignment based on cell type composition and gene expression between oligodendrocytes and astrocytes (top) with alignment similarity between the two cell type trajectories across the four time points (bottom). (F) Single gene trends confirm BMP and NOTCH pathways upregulation in astrocyte trajectory development that strongly correlates with steadily increased and sustained NFIA expression.

### Single cell RNA sequencing reveals robust oligodendrocyte lineage induction with minimal astrocyte lineage contamination following NOTCHi

The overall datasets included in our analysis involve four time points (day 12, day 22, day 32, and day 43). The first two time-points are identical across the three conditions (*Base D12 – Base D22*). The day 32 time point involves two distinct treatment paradigms: BMPi starting at day 26 (*Base D32*) versus combined BMPi and NOTCHi starting at day 26 (*Mature D32*). The day 43 time point involves three distinct treatment paradigms: BMPi only, from day 26 - 43 (*Mixed glial D43*), both BMPi and NOTCHi from day 26 - 43 (*Mature D43*), and BMPi from day 26 - 43, but delayed NOTCHi from day 32 - 43 (*expansion D43*) (**Fig. 6A**). We combined all the data across the four time points and treatment conditions in one single UMAP and included curated annotations of cell types based on canonical markers (**Fig. 6A, S6A-D**). Single gene projection UMAPs show a developmental characterization of oligodendrocyte lineage progression across the three protocols based on markers of pMN domain, glia-competent state, OPC identity, oligo-commitment, and mature and myelinating oligodendrocytes (**Fig. 6B**). Quantification of cell types based on the single cell data shows approximately 40% astrocytes in *mixed-glial D43*, while both *Expansion D43* and *Mature D43* showed an almost complete absence of astrocytes (**Fig. 6C, S6C**). On the other hand, oligodendrocytes represented approximately 65% of the total cells in *mature day 43* (**Fig. 6C, S6D**) similar to our flow-based quantification. The oligodendrocyte population represented around 30% in the *Expansion D43* and around 20% in the *Mixed glial D43* condition. The percentage of OPCs was highest in *Expansion D43* (28%) (**Fig. 6C**) but also enriched in *Mature D32*, supporting the hypothesis that a shorter time period of NOTCHi, rather than a specific delay of NOTCHi is critical for enriching *PDGFRα* expressing OPC-like progenitors (**Fig. 6D, E**). To further illustrate the relative enrichment of lineage specific transcriptional profiles, we separated the conditions that received BMPi plus NOTCHi (both at day 26 and day 32) versus those that received BMPi only. Cells corresponding to *Expansion D43* and *Mature D32* conditions were closely matched in transcriptional profiles further highlighting the role of the short-term NOTCHi paradigm in triggering OPC-like state and suggesting that both D26 and D32 cells are similarly competent to respond to NOTCHi. On the other hand, *Mixed-glial D43* and *Mature D43* show highly distinct UMAP patterns (**Fig. 6D**) enriched for either AQP4, GFAP-positive astrocytes versus mature oligodendrocyte markers including MBP and MOG, respectively. The *Expansion D43* and *Mature D32* clusters are enriched for PDGFRα and SOX10 expression corresponding to the OPC and immature oligodendrocyte lineage (**Fig. 6E**).

**Figure 6.**
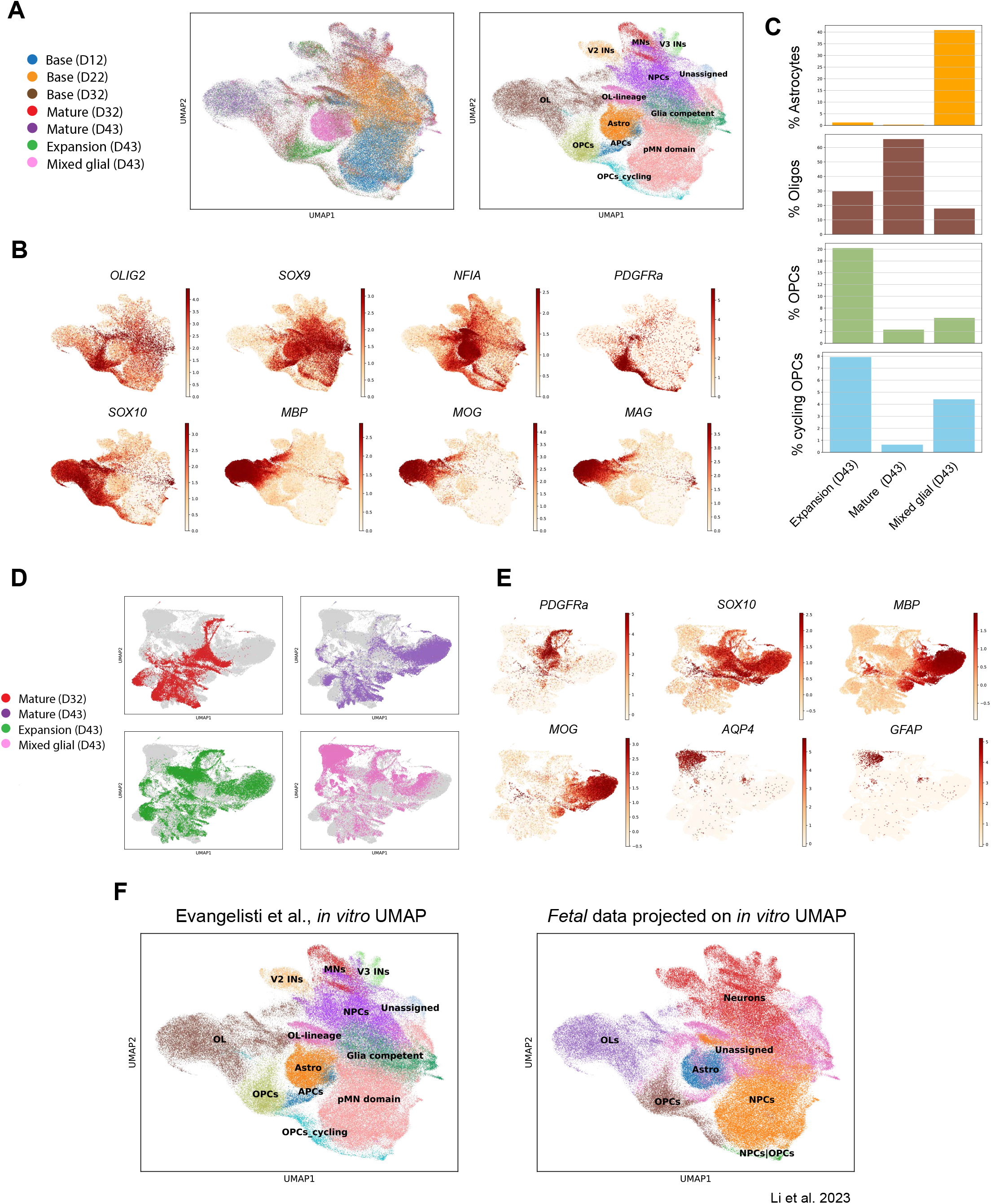
Single cell RNA sequencing reveals robust oligodendrocyte lineage induction with minimal astrocyte lineage contamination following NOTCHi. (A) Concatenated UMAP shows all three protocols (mixed-glial, expansion and mature) at day 12-22-32-43. Left: three protocols across four time points. Right: corresponding curated cell type annotations. (B) Gene expression plots describe the transition from glia competent cells (OLIG2-NFIA-SOX9) to oligo progenitors (PDGFRα-SOX10), to mature oligodendrocytes (MBP, MOG, MAG). (C) Quantification of astrocytes, oligodendrocytes, OPCs across all three protocols at day 43. (D) UMAP plots illustrating the contribution of cells from each differentiation protocol. (E) Gene expression plots show canonical markers that characterize OPCs, mature oligodendrocytes and astrocytes. (F) Left: cell type annotations of our in vitro data. Right: Majority voting (probability = 0.5) from CellTypist shows a close match of the oligodendrocyte lineage between our in vitro data and a human fetal spinal cord atlas^48^.

Finally, we compared our single cell data with a publicly available human fetal spinal cord atlas^48^. We used *CellTypist*^49^ to train a model for cell type discrimination based on the *in vivo* dataset. To enable accurate identification of glial populations, we refined the original published annotations, which did not include astrocytes. This refinement incorporated well-established marker genes, allowing us to distinguish both oligodendrocytes and astrocytes, with astrocytes identified based on canonical markers such as *GFAP* and *AQP4* (**Fig. S6F**).

We found that the hPSC-derived oligodendrocytes and astrocytes faithfully matched the human fetal cell counterparts with a probability threshold greater than 0.5 (**Fig 6F, S6E**), showing that both lineage cells faithfully matched the corresponding cell types in the *in vivo* fetal cell atlas. However, we found that, the earlier time points (pMN domain and glia competent stage) were not properly recapitulated in the fetal data. This was likely due to the fact that these early hPSC-derived lineages *in vitro* reflect an earlier developmental stage that is not properly captured in the corresponding *in vivo* fetal atlas of 5 - 12 weeks p.c. reported in the published study ^48^ (**Fig. 6F**).

### Cells show signs of myelination as early as day 42 both in 2D and 3D (native) environments

One of the major challenges of current protocols is to achieve evidence of myelination in hPSC-based systems without requiring many months of *in vitro* differentiation or *in vivo* transplantation. As our differentiation paradigm involves a 3D-organoid-type differentiation paradigm, we first assessed whether in this native environment, cells show MBP expression along NF-H neuronal axons present within these cultures (**Fig. 7A**). Next, we dissociated organoids at day 32 and replated them in a 2D culture format for further cell characterization. We allowed further differentiation *in vitro* to day 42 in *Mature* and *Expansion* conditions and showed that cells were ramified, positive for mature markers such as MBP and MOG, and aligned along neural axons (**Fig. 7B, S7A**). In a follow-up study, we kept the cells in the organoid environment until day 42, followed by dissociation and plating in 2D culture followed by analysis at day 52. We observed a further increase in morphological complexity and neuronal interactions (**Fig. 7C, S7B**). These studies were based on keeping cells within their native system where neurons (mostly of motor neuron identity) were found as a differentiation side-product (**Fig. S6B**). However, to further test the utility of our culture platform, we developed conditions for cryopreservation to test their myelination capacity upon thawing of cryopreserved populations in heterologous culture systems. We sorted SOX10+ cells at day 32 using our double reporter line and immediately froze them using a controlled rate freezer. In parallel, we differentiated a doxycycline inducible iNGN2 hPSC line to rapidly generate cortical-like neurons. Around day 15 of neuronal differentiation, we added sorted SOX10+ cells on top of 2D plated iNGN2 neurons and maintained these co-cultures for 15 additional days. We observed a further increase in the morphological complexity of the oligodendrocytes with a single cell contacting numerous axons (**Fig. 7D**, top panels). Importantly, axons (NF-H+) closely overlapped with MOG and MBP fibers, forming extended axonal structures surrounded by oligodendrocytes mimicking early steps of myelination in our culture system (**Fig. 7D**, lower panels).

**Figure 7.**
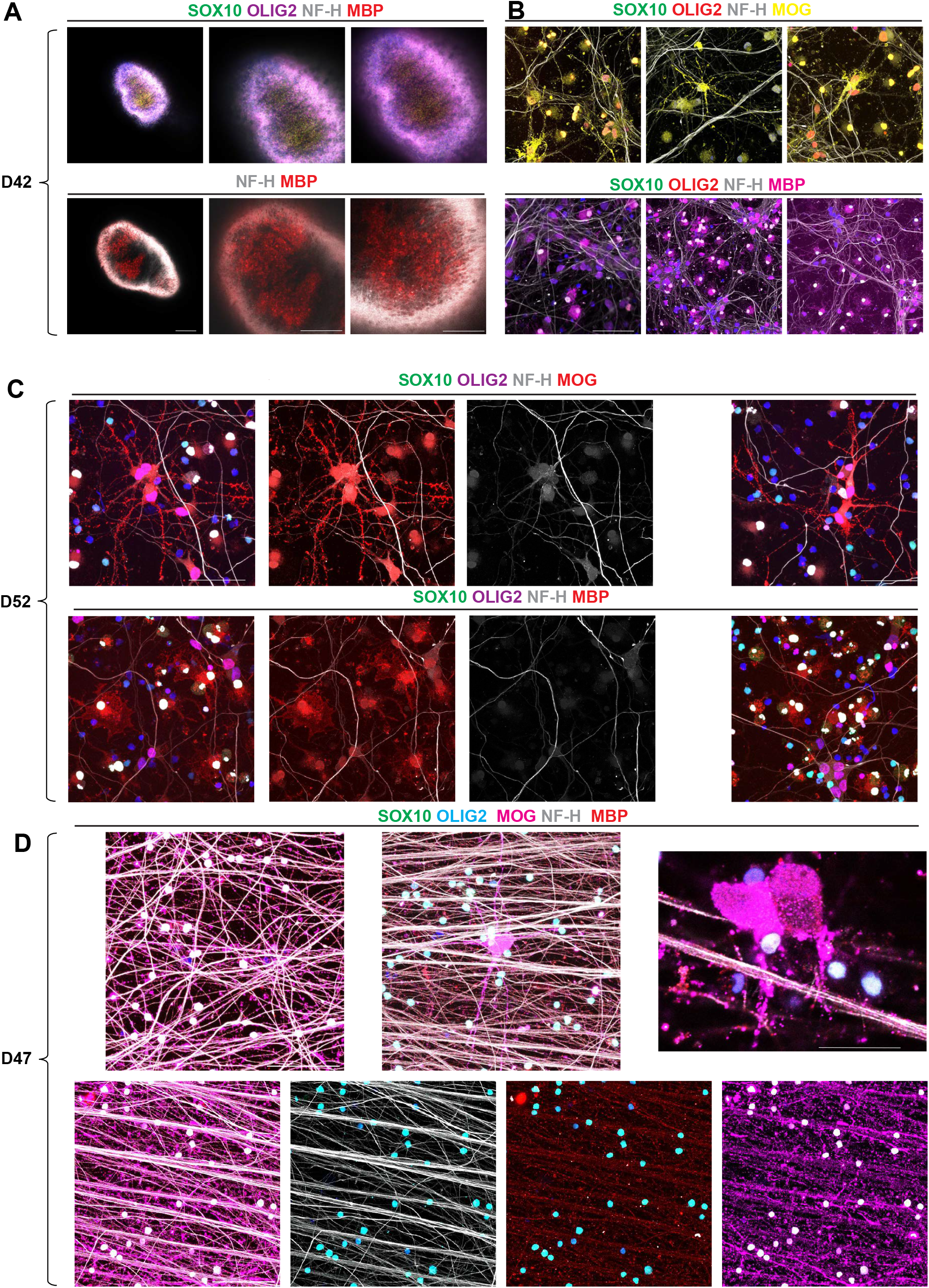
Cells show signs of myelination as early as day 42 both in 2D and 3D (native) environments. (A) Representative confocal images show at day 42, *MBP* staining widely spread across the whole organoid while most of the cells remain *SOX10* and *OLIG2* positive (from left to right - 10X and 20X magnification). (B) Cells dissociated at day 32, plated and cultured in 2D and fixed at day 42; we find widespread oligodendrocytes positive for *MOG* and *MBP* along *NF-H* positive axonal neurons (upper panel at 63X, lower panel at 40X). (C) At day 52, the oligodendrocytes appear to have more complex structures, and the glial processes reach further distances alongside neurons (63X). (D) Co-culture for 15 days of previously sorted *SOX10* positive cells (at day 32) with iNGN2-derived neurons and fixed at day 47, shows an intricate network of myelin (MOG and MBP) that runs along *NF-H* positive axons. Scale bar: 100µm.

## Discussion

Over the last few years, several hPSC-derived CNS lineages have progressed towards translational applications including in human disease modeling and drug screening^50–55^, as well as in cell therapy^56–58^. However, the derivation of oligodendrocytes has remained challenging due to the protracted timelines of differentiation^20,21,59^ and the often heterogeneous cultures that emerge from these protocols, which complicate their use in translational applications including clinical grade manufacturing.

Our study focused on oligodendrocyte differentiation using a spinal cord differentiation paradigm, similar to past protocols^19^. In fetal development, the first wave of oligodendrocyte development occurs in the spinal cord, particularly the ventral compartment, and therefore we hypothesized it might be this domain that allows for the derivation of oligodendrocytes most quickly. Nevertheless, it will be important in future studies to confirm that our protocol involving BMPi and NOTCHi similarly applies to oligodendrocytes developing in other brain regions including forebrain oligodendrocytes and those that arise from more dorsal compartments. We adopted an organoid platform for oligodendrocyte induction, as it may better recapitulate *in vivo* environments and architecture, which include extensive cell-to-cell interactions^60–62^ and neuronal activity, which is thought to be critical for oligodendrocyte development and maturation^4,63,64^.

A key finding of our study is the identification of signaling pathways that act on bipotent, glial competent precursors emerging from the pMN domain to control astrocytic versus oligodendrocyte lineage choice and to promote oligodendrocyte differentiation and maturation. We have identified BMP and Notch signaling as critical regulators of these fate choices, and we identified dual inhibition of these pathways as an efficient strategy for the rapid and efficient differentiation of hPSC-derived oligodendrocytes. Previous work has shown a role for BMP signaling in promoting astrocyte differentiation, while inhibiting neurogenesis^40^. BMP activation was proposed as a major suppressor of oligodendroglia development of glia-restricted bipotent progenitor cells^65^. One possible mechanism involves the BMP-mediated promotion of OLIG1 and OLIG2 translocation to the cytoplasm, therefore preventing the transcription of OLIG1/2-dependent genes involved in oligodendrocyte differentiation^41^. Furthermore, inhibition of the Notch signaling pathway has been described to inhibit primary astrocytes development^66^, while its activation can inhibit oligodendrocyte differentiation^67^ and induce astrogliogenesis^68^. Our differentiation protocol involved an organoid-based culture system. However, preliminary data suggest that dual inhibition of NOTCH and BMP signaling is similarly effective under 2D oligodendrocyte differentiation conditions.

We made additional changes to the overall differentiation protocol by using RA receptor agonists rather than all trans retinoic acid to activate retinoid signaling, which resulted in a more robust induction of pMN lineage markers and reduction of more ventral fates including FOXA2 expression. Activation of retinoic acid signaling is thought to maintain cells within the pMN domain working against the opposing gradient of SHH coming from the notochord (**Fig. 2A**); However, the mechanism of differential action comparing RA agonist versus ATRA treatment remains to be determined; whether RA agonists simply improve overall potency of pathway activation and avoid major off-target effects, or whether the specific balance of RAR versus RXR activation is critical. Similarly, we observed improved potency of FGF18 in maintaining OLIG2 expression and the bipotent precursor stage compared with no FGF or FGF2 treatment, but we have not tested all the possible FGF ligands or levels of receptor subtype activation to determine qualitatively unique patterns of FGFR activation versus quantitative differences between FGF18 and FGF2. Finally, we used cyclopamine, a strong SHH signaling inhibitor, to suppress the small percentage of contaminating FOXA2+ ventral floor-plate like cells, which may be biased towards astrocytic differentiation.

Our study used scRNAseq and trajectory analysis to demonstrate a tight temporal link between BMP and Notch signaling pathways in the glial differentiation trajectories with elevated levels of pathway activation leading to astrocytic fates. These data are compatible with previous work showing that the transition from oscillatory Notch signaling to constant high levels of activation marks the transition from a neural stem cell to an astrocyte committed fate^69,70^, and we further validated this link by demonstrating efficient BMP4-induced astrocyte induction. Conversely, we found that inhibition of BMP is critical to reduce the differentiation of bi-potent progenitor cells into astrocytes and to preserve oligodendrocyte differentiation potential. We observed two main effects of NOTCHi: early inhibition promoted the conversion of bi-potent precursors into committed pre-myelinating oligodendrocytes and sustained inhibition acted as a maturation switch promoting terminal differentiation of oligodendrocytes. While NOTCHi is a well-known strategy to promote differentiation of neurons from neural precursor cells^45^, its role in oligodendrocyte differentiation is less clear. Nevertheless, NOTCH inhibition has been proposed as a strategy to promote oligodendrocyte differentiation and myelination in primary OPCs^71^ including OPCs in the spinal cord^72^, and in classic rat optic nerve studies^67^. These findings are compatible with the effect reported here on terminal differentiation and maturation of oligodendrocyte lineages. The early role of NOTCHi in driving bipotent precursor cells towards OPC induction was highly robust in our study, but is less well understood mechanistically, as some *in vivo* studies report that a positive regulation of Notch signaling in OPC induction such as in the zebrafish where Delta-Notch signaling is required for spinal cord oligodendrocyte specification^73^.

Our modular strategy to derive three distinct glial cell products should enable a broad set of applications in translational medicine. The application for the *mature* oligodendrocyte protocol (sustained NOTCHi) may involve myelination-related assays and drug-based screening efforts at promoting myelination. In contrast, the expansion condition (short NOTCHi), may yield an expandable SOX10/PDGFRα progenitors suitable for future cell therapy efforts, where immature and cycling, yet committed, cells may enable engraftment and widespread migration as compared with O4+ non-cycling oligodendrocytes. Finally, the *mixed-glial* (*no NOTCHi*) protocol gives rise to a heterogeneous population of both oligodendrocytes and astrocytes. Such a mixed glial population could serve as a source of both astrocytes and oligodendrocytes for use in triculture with neurons or in tetra-culture paradigms with neurons and microglia for human disease modeling. One main advantage of the mixed glial protocol is the rapid derivation of oligodendrocytes but also of AQP4+ astrocytes. AQP4 is one of the most specific markers for bona fide astrocytes^74^ and not robustly expressed in most current rapid astrocyte differentiation protocols^75^.

We developed a simple, fast, and reliable protocol using defined extrinsic growth factors and small molecules to generate oligodendrocytes without the need of any transgene expression. The study should facilitate the application of human oligodendrocytes in the broader stem cell and neuroscience community. In particular, the work may contribute to the development of novel drug and cell-based strategies for the treatment of a broad range of demyelinating disorders such as multiple sclerosis, multiple systems atrophy (MSA), genetic leukodystrophies, radiation/chemotherapy-induced demyelination, which can lead to significant and debilitating cognitive impairment, and age-related demyelination of the CNS.

## Limitations

While we showed rapid and efficient generation of mature oligodendrocytes, we cannot conclusively state that the cells rapidly myelinate neurons in our culture system. We demonstrate the proximity of MOG/MBP myelin fibers to NF-H axons, but preliminary electron microscopy images at these early differentiation stages did not provide conclusive evidence of myelin wrapping^76^, and future *in vivo* transplantation studies may be required to assess the full potential of these cells for *myelination*.

## Data Availability

The single-cell RNA sequencing (scRNA-seq) datasets generated in this study have been deposited in ArrayExpress via Annotare under accession number E-MTAB-16931. The data will be made publicly available upon publication of the peer-reviewed manuscript.

## Acknowledgements

We thank members of the Studer laboratory for helpful discussions. We thank Elizabeth Calder for the specific advice on use of the RAR and RXR agonists. We thank Ronan Chaligne and the Single Cell Analytics Innovation Lab (SAIL) team at MSK for their assistance with single cell library preparation and their valuable advice on transcriptomic approaches and cell hashtag multiplexing. We thank the Integrated Genomic Operation (IGO), the Molecular Cytology Core Facility and the Light Microscopy Instrument Cluster at MSKCC. This project was supported in part by grants from the National Institute of Mental Health R01MH135403, the National Institute of Neurological Disorders and Stroke R01NS128087 and by center grant support from the National Cancer Institute P30CA008748. A.E. was supported in part by Dompe fellowship, V.D.B by Kravis Wise Fellowship.

## Author Contributions

L.S. and A.E. conceived and designed experiments, analyzed and interpreted data, and wrote the manuscript. A.E. performed organoid differentiation, imaging experiments, flow cytometry experiments, and single cell RNA-seq analysis. S.P. contributed to the early conceptualization, study design and interpretation of the data. J.J. provided expertise on early protocol development and contributed to the study design. R.M.W. provided technical support on scRNAseq sample preparation and expertise on organoid differentiation. V.D.B.: provided expertise and feedback on single cell analyses. Y.W. and T.Z. generated the SOX10-GFP/OLIG2-tdTomato dual reporter cell line use throughout the study.

## Declaration of interests

L.S. is a co-founder, scientific advisor, and has received research support from BlueRock Therapeutics. L.S. is a co-founder and scientific advisor of DaCapo Brain Science. L.S. and A.E. are inventors of a patent application owned and filed by Memorial Sloan Kettering Cancer Center, based on the differentiation technologies presented here.

## Methods

### Cell lines

Human pluripotent stem cells [hPSCs; WA09 (H9; 46XX), WA01 (H1; 46XY), SOX10-GFP-OLIG2-tdTomato-H2B in H9 background, and MSKSRF01] were grown on Vitronectin (VTN-N, Thermo Fisher #A14700) coated 6-well plate with Essential 8 media (Life Technologies #A1517001) supplemented with tankyrase inhibitor, XAV (#N/A). hPSCs were passaged every 3-4 days by EDTA, and passage 5-40 hPSCs were used for the experiments. The H9 SOX10-GFP/OLIG2-tdTomato dual reporter line was generated by sequential CRISPR/Cas9-mediated homology-directed repair (HDR), as previously reported^28^ (**Supplementary Fig. 1A**). The OLIG2-tdTomato reporter was introduced into the validated SOX10-GFP reporter line^77^ by electroporation of an sgRNA targeting the OLIG2 locus together with a donor plasmid containing a P2A-H2B-tdTomato cassette followed by a floxed neomycin resistance cassette (loxP-PGK-Neo-loxP).

Neomycin (G418) selection was applied, and correctly targeted double knock-in clones were identified by PCR and Sanger sequencing. Drug selection cassettes were removed by Cre-mediated recombination via electroporation of a Cre-expressing plasmid (1 µg) into a confirmed double knock-in clone. Successful excision was verified by PCR and Sanger sequencing (**Supplementary Fig. 1B**). Expression of both reporter alleles was further validated using CRISPRa-mediated gene activation, as previously described^28,78^. Briefly, the SAM-TET1 system and sgRNAs targeting SOX10 and OLIG2 were electroporated into the dual reporter line to transiently activate endogenous gene expression. GFP and tdTomato fluorescence was detected 48 hours post-electroporation (**Suppl. Fig. 1C**).

### Directed differentiation into spinal cord oligodendrocytes (OLs)

hPSCs were dissociated into smaller aggregates using EDTA for 5 minutes at RT and plated at 10K cells/well into 96V-bottom low attachment dishes (S BIO PrimeSurface #MS-9096VZ) in Essential 6 media (Life Technologies #A1516401) 10µM ROCK inhibitor (Y-27632, R&D systems #1254). The plate was then spun down at 1100 rpm for 2 minutes concluding day −1 of differentiation. On the following day, the media was replaced with Neurobasal (Life Technologies)/N2(Stem Cell Technologies)/B27(Life Technologies) media containing 2mM L-glutamine, 250nM LDN (Stemgent # 04-0074-02), 10μM SB431542 (R&D systems #1614), 3μM CHIR99021 (R&D systems #4432) and 100μM AA (Sigma #4034-100g), and change every day until day 6. On day 6, newly formed organoids were transferred from the 96V bottom plate onto a 10cm dish and cultured in previous media with the addition of 2μM SAG (Selleck #S6384), 0.2μM RXR agonist (Tocris Bioscience #5920) and 0.25μM RAR agonist (Tocris Bioscience #3505). The media was changed every day until day 12. On day 12, the media was changed to Neurobasal/B27/L-Glu supplemented with 20ng/ml BDNF (R&D #248-BD), 20ng/ml GDNF (Peprotech # 450-10),100μM AA (Sigma #4034), 50ng/mL FGF18 (Peprotech #100-28), 20ng/ml PDGF-AA (R&D #221-AA-100), 2μM SAG (Selleck #S6384), 0.2μM RXR agonist (Tocris Bioscience #5920) and 0.25μM RAR agonist (Tocris Bioscience #3505). The medium was changed every other day until day 22. On day 16-18, organoids were chopped into 4 small pieces using a precision needle (BD #305106) and the equivalent of one 10cm dish organoids were transferred among two new dishes in the same media as described before. On day 22, GDNF, SAG, RXR and RAR agonists were withdrawn, and 5μM cyclopamine (Selleck #S1146) was added to the mix of small molecules. The media was changed every other day until day 26. On day 26, 2µM DMH1 (Tocris Bioscience #4126) and 10μM DAPT (R&D #2634) were added to the media until day 32, with media change every other day. On day 32, the media was changed to Neurobasal/B27/L-Glu supplemented with 20ng/ml BDNF (R&D #248-BD), 100μM AA (Sigma #4034), 2µM DMH1 (Tocris Bioscience #4126) and 10μM DAPT (R&D #2634) until day 42 and onwards with half media change every 2-3 days. On day 42, organoids were dissociated for 1h using the Papain Dissociation System (Worthington Biochemical Corporation #LK003150) and used for any downstream application and/or QC by 1. replating single cells in a 96-well plate at a density of 300k cells/cm^2^ in Neurobasal/B27/L-Glu supplemented with 20ng/ml BDNF (R&D #248-BD), 100μM AA (Sigma #4034), 2µM DMH1 (Tocris Bioscience #4126), 10μM DAPT (R&D #2634) and 10µM ROCK inhibitor (Y-27632, R&D systems #1254), or 2. FACS analyzed O4 positive population (see below for directions).

### Immunohistochemistry

Cells were fixed in 4% paraformaldehyde (PFA) (Affymetrix #MFCD00133991) in DPBS for 12 min at room temperature. Cells were subsequently washed with DPBS++. Then samples were permeabilized with 0.1% Triton X-100 and blocked with 10%NGS + 1%BSA + 0.1% Triton X-100 in DPBS++. The samples were subsequently incubated with primary antibody in 10%NGS/1%BSA/0.1%TritonX-100 (DPBS++) overnight at 4°C with gentle rocking. The next day, after washing with DPBS, the samples were incubated with secondary antibody conjugated with Alexa Fluor 488-555-647-, or −750 (Thermo Fisher) diluted at 1:400 in 10%NGS/1%BSA/0.1%TritonX-100 (DPBS++) for 30 minutes at room temperature with gentle rocking. Then the samples were washed with DPBS++ and counterstained with 41, 6-diamidino-2-phenylindole (DAPI) (Sigma, #D9542). Images were taken using a Zeiss inverted fluorescence and a Leica STELLARIS confocal microscope. Goat anti-FOXA2 (1:200, R&D), rat anti-MBP, rat anti-PLP1 (1:100), rabbit anti-MOG (1:300), mouse anti-SOX9 (1:100), mouse anti-SOX10 (1:100), rabbit anti-OLIG2 (1:500), mouse anti-OLIG2 (1:100), mouse anti-NKX2_2 (1:100), rabbit anti-NFIA (1:500), chicken anti-NFH (1:500), rabbit anti-GFAP (1:2000), mouse anti-ISLET1 (1:100) were used for immuno-fluorescent staining. Goat and donkey anti-mouse, goat, rabbit or chicken secondary antibodies conjugated with Alexa Fluor-488, Alexa Fluor-555, Alexa Fluor-647, or Alexa Fluor-750 fluorophore (1:400, Life technologies) were used. Nuclei were counterstained by DAPI.

### Flow Cytometry

Organoids were dissociated for 1h using the Papain Dissociation System (Worthington Biochemical Corporation #LK003150). Single cell suspensions were stained in FACS buffer (DMEM/F12 + 2%FBS+ 0.2µM EDTA + ROCK inhibitor) for 45 minutes on ice with O4-conjugated-APC/PE, or O1-conjugated-APC antibodies at a concentration of 10μL and 50μL per 100K cells in 100μL FACS buffer, respectively. Cells were washed three times in FACS buffer and then counterstained with 4⍰, 6-diamidino-2-phenylindole (DAPI) (Sigma, #D9542). Cells were passed through a 30μm filter (Sysmex #04-004-2326) and analyzed using Cytek Aurora Flow Cytometer.

### Fluorescence-Activated Cell Sorting (FACS)

For cell sorting, (SOX10 at day 32 and O4 at day 42) organoids were dissociated for 60 minutes using the Papain Dissociation System (Worthington Biochemical Corporation #LK003150). Cells were washed three times in FACS buffer and then counterstained with 4⍰, 6-diamidino-2-phenylindole (DAPI) (Sigma, #D9542). Cells were filtered through a 30μm strainer (Sysmex #04-004-2326), sorted them in the FACS buffer using BD FACSymphony S6 or FACSDiscovery S8 cell sorters, collected them in 5mL polystyrene tubes. Sorted cells were spun down and washed twice in Neurobasal/B27/L-Glu media supplemented with 20ng/ml BDNF (R&D #248-BD), 100μM AA (Sigma #4034), 2µM DMH1 (Tocris Bioscience #4126), 10μM DAPT (R&D #2634) and 10µM ROCK inhibitor (Y-27632, R&D systems #1254). After counting, cells were plated for different downstream assays or frozen at a cell density of 500k cells/mL of STEM-CELLBANKER. Controlled rate freezer (ThermoFisher) was used to cryopreserve cell product.

### *In vitro* myelination assay

iNGN2 neurons were generated using a published protocol in μ-Plate 96 Well Square (Ibidi #89626) for high-resolution microscopy. At 9-10 days of neuronal differentiation, as neurites extended throughout the culture, SOX10+ oligodendrocytes, previously frozen, were thawed on top of forming neurons at a concentration of 180k cells/cm^2^ in Neurobasal/B27/L-Glu media supplemented with 20ng/ml BDNF (R&D #248-BD), 100μM AA (Sigma #4034), 2µM DMH1 (Tocris Bioscience #4126), 10μM DAPT (R&D #2634) and 10µM ROCK inhibitor (Y-27632, R&D systems #1254). Media was half-changed every 2-3 days without ROCK inhibitor. After 15 days, cells were washed with 1×PBS and fixed in 4% paraformaldehyde.

### Single Cell RNA Sequencing

Organoids were washed once in 1×PBS and enzymatically dissociated using Papain Kit (Worthington, #LK003150) at 37 °C and 5% CO_2_ for 60 min on an orbital shaker. Following enzymatic digestion, organoids were mechanically dissociated into single cells by repetitive pipetting using a 1,000 µL pipette tip. Papain activity was quenched, and cells were washed according to the manufacturer’s instructions. For sample multiplexing, two to four samples per time point were stained separately with Cell Hashing antibodies. Briefly, single-cell suspensions were incubated with human TruStain FcX (BioLegend, #422301) for 10 min at room temperature to block Fc receptors, followed by staining with TotalSeq™ Cell Hashing antibodies (e.g., TotalSeq™-A0251 anti-human Hashtag 1 antibody, BioLegend, #39460) for 30 min on ice. Cells were washed three times with Cell Staining Buffer (BioLegend, #420201), passed through a 30 µm cell strainer, and counted using trypan blue exclusion on a Countess II Automated Cell Counter (Thermo Fisher Scientific). Cells were then pooled into a single low-protein-binding 1.5 mL microcentrifuge tube and resuspended in 1× PBS containing 0.04% BSA. After quality control, 30,000–60,000 cells were loaded onto a Chromium Next GEM Chip G (10x Genomics, #1000120). Gel bead-in-emulsion (GEM) generation, reverse transcription, cDNA amplification, and library preparation were performed using the Chromium Next GEM Single Cell 3⍰ Reagent Kit v3.1 (10x Genomics, #1000268) according to the manufacturer’s protocol. cDNA amplification was performed using 11–12 PCR cycles, and 30–240 ng of amplified cDNA was used for library construction with 10–16 PCR cycles. Indexed libraries were pooled at equimolar concentrations and sequenced on an Illumina NovaSeq 6000 platform using paired-end 28/88 bp reads with NovaSeq S1 or S4 reagent kits (100- or 200-cycle).

### Single Cell Data analysis

Raw sequencing data were demultiplexed and aligned to the human GRCh38 reference genome using Cell Ranger (v6.1.2). Count matrices were imported into Python (v3.12) for downstream analysis using *Scanpy* within Jupyter Notebook (v7.0.8). Cells were retained for downstream analyses if they expressed at least 500 genes and contained fewer than 10% mitochondrial gene-derived counts. Genes expressed in fewer than three cells were excluded. Additional quality control filters were applied on a per-dataset basis to remove low-quality cells and technical artifacts. Gene expression values were normalized on a per-cell basis, log-transformed, and scaled without explicit selection of highly variable genes. Principal component analysis (PCA) was performed using the full gene expression space, and the resulting components were used for downstream neighborhood graph construction. To correct for batch effects across samples, time points, or experimental conditions, batch-balanced k-nearest neighbors (BBKNN) integration was applied where indicated. Integrated neighbor graphs were used for dimensionality reduction and visualization using Uniform Manifold Approximation and Projection (UMAP). Clustering was performed using the Leiden algorithm with resolution parameters adjusted depending on the biological question being addressed. Trajectory inference was performed for selected analyses using *Palantir* to model differentiation dynamics and pseudo-temporal ordering of cells. For these analyses, *Palantir* was applied to integrated or non-integrated embeddings as appropriate, and lineage progression was inferred based on diffusion maps and probabilistic fate assignment. Gene regulatory dynamics and temporal gene–gene relationships were further explored using *genes2genes*, enabling the identification of transcriptional programs associated with lineage transitions and maturation states. As a final comparative analysis, *CellTypist* was used as a supervised machine-learning framework that assigns probabilistic cell-type labels by projecting query single-cell transcriptomes onto reference datasets. This approach was applied to compare *in vitro* derived cell populations with published human fetal single-cell reference data. Cell-type assignments were considered when the predicted class probability was ≥ 0.5 and were not used for primary cell type annotation.

### Quantification and statistical analysis

Quantification of immunofluorescence images was performed manually using FIJI (ImageJ). Statistical analyses were conducted using GraphPad Prism 10. For statistical analysis experiments were performed with at least three independent biological replicates derived from independent differentiations. Data are presented as mean ± standard error of the mean (SEM). Statistical significance is indicated as follows: *P* < 0.05, *P* < 0.01, *P* < 0.001, *P* < 0.0001. Flow cytometry data were analyzed and visualized using FlowJo (v10). Single-cell analyses were performed and visualized using Scanpy in Jupyter Notebook (v7.0.8).

## Legends to Supplementary figures

**Supplementary Fig 1.**
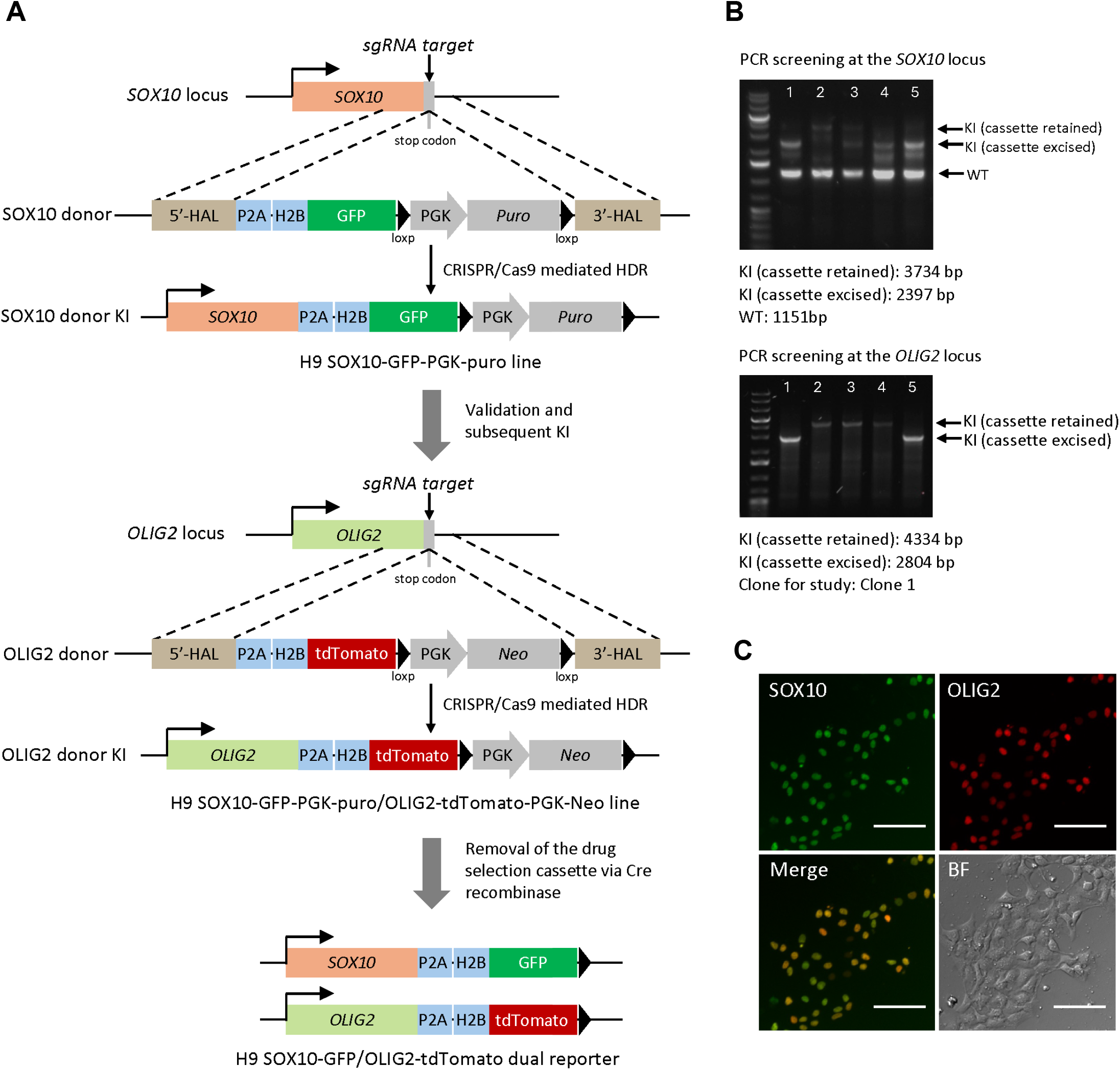
Generation and verification of the H9 SOX10-GFP/OLIG2-tdTomato dual reporter. (A) Schematic illustration of the sequential generation of the H9 SOX10-GFP/OLIG2-tdTomato dual reporter line. (B) PCR verification of dual reporter clones following Cre-mediated excision of the drug selection cassettes. (C) Fluorescence images of H9 SOX10-GFP/OLIG2-tdTomato cells 48 h after electroporation with the SAM-TET1 system and sgRNAs targeting SOX10 and OLIG2. tdTomato (OLIG2), GFP (SOX10), and merged images are shown. BF, bright field. Scale bar, 10 μm.

**Supplementary Fig 2.**
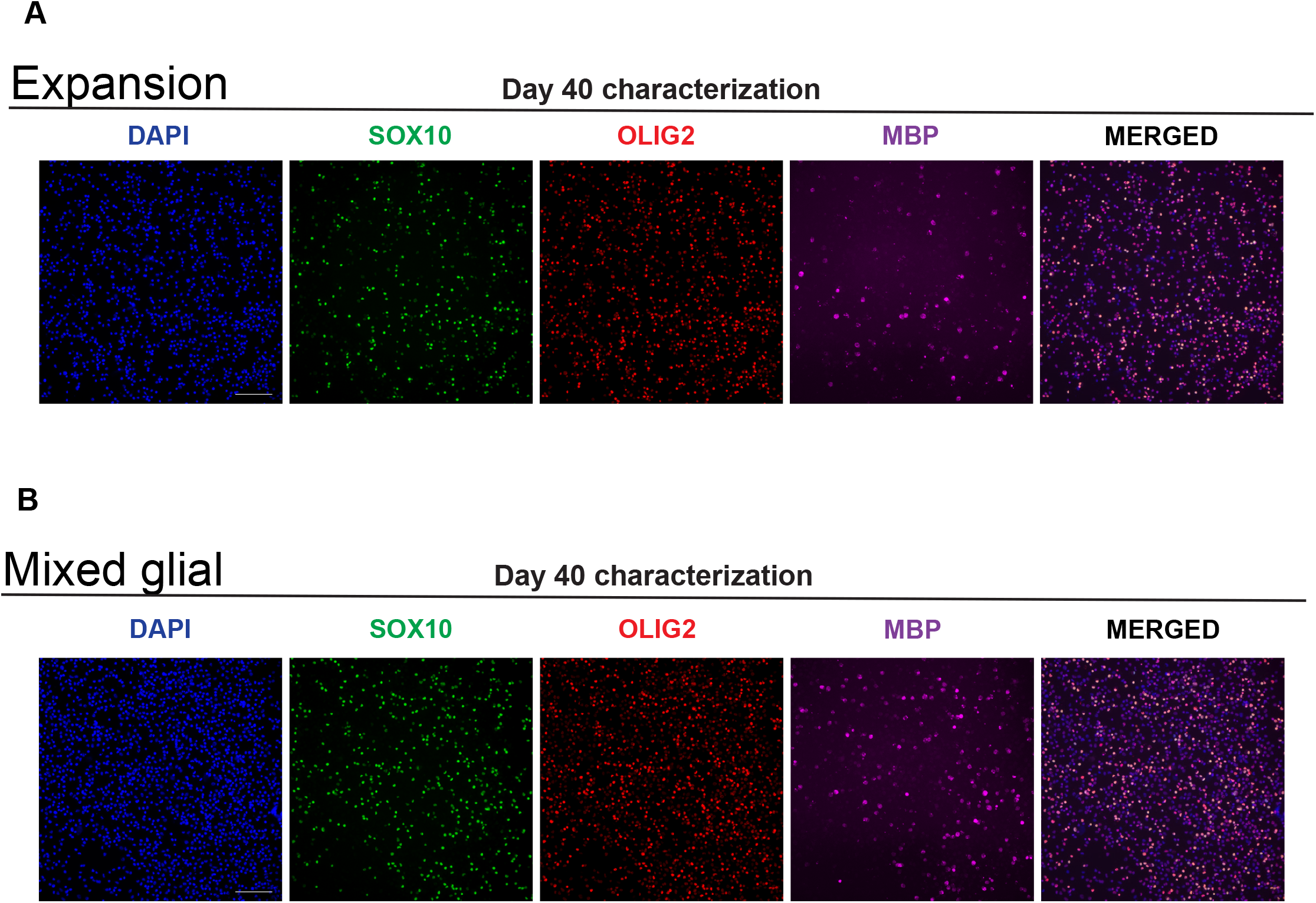
Validation of oligodendrocyte identity across the three differentiation protocols. (A) Representative immunofluorescence images show cells identity at day 42 of the (A) *expansion* and (B) *mixed-glial* protocols, mostly positive for OLIG2, SOX10, and myelin basic protein (MBP). Scale bar: 100μm.

**Supplementary Fig 3.**
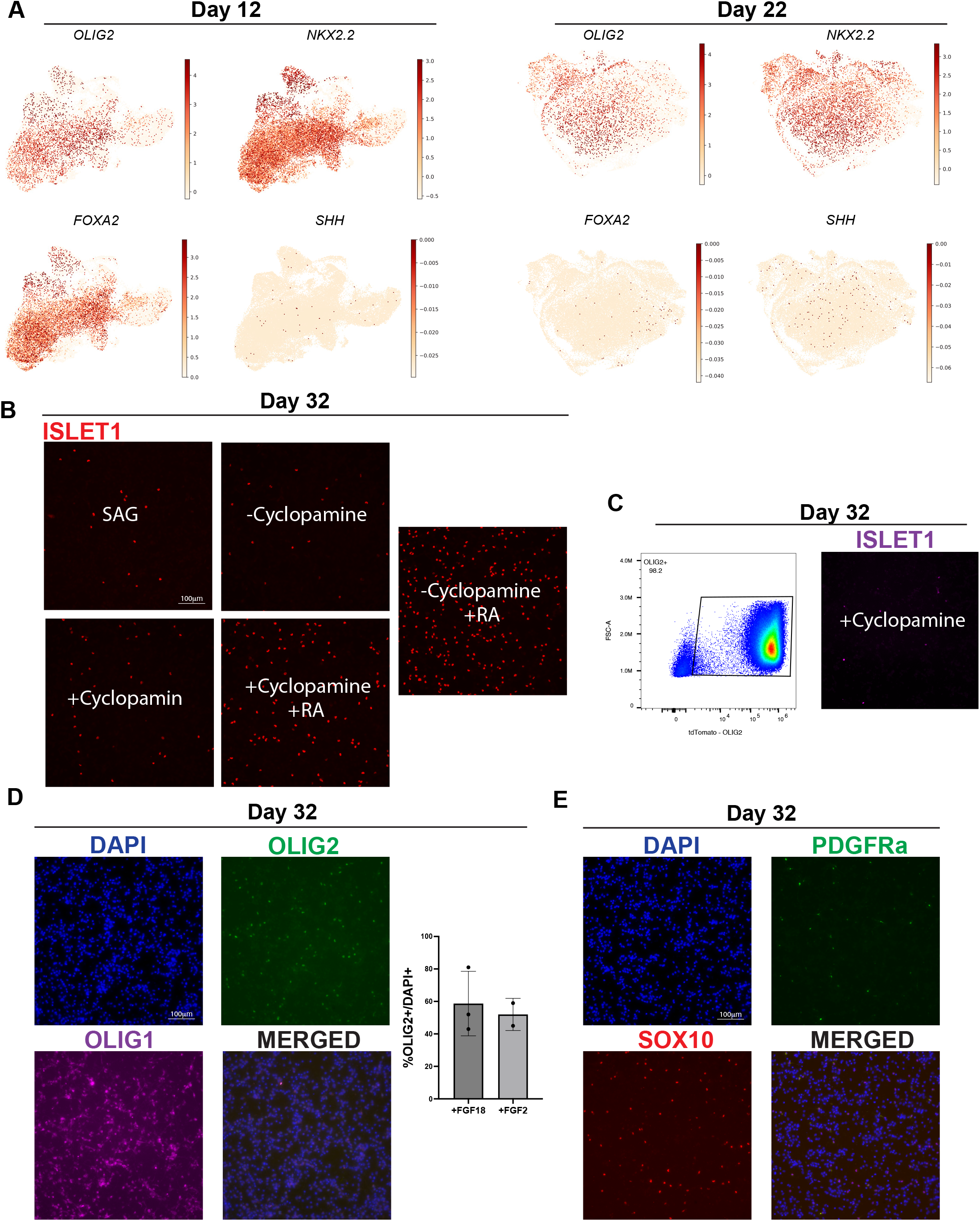
RA agonists, FGF18 and SHH-inhibitor keep cells within the pMN domain limiting FOXA2 floor-plate cells proliferation and motor neurons development. (A) Strong expression of *OLIG2-NKX2-2* at day 12 and an intermediate progenitor *FOXA2* population that does not express common markers of floor plate cells such as SHH (left). By day 22, single gene expression UMAPs show a complete FOXA2 population suppression while retaining pMN domain identity. (B) The inhibition of endogenous SHH, by cyclopamine, and the removal of retinoic acid agonists limit *ISLET1+* motor neurons development, while keeping high *OLIG2* throughout the differentiation (C). (D) By day 22, cells are strongly positive for *OLIG2* and *OLIG1*, and by day 32, some of these cells transition from OPCs to SOX10 committed oligodendrocytes (E). Scale bar: 100μm.

**Supplementary Fig 4.**
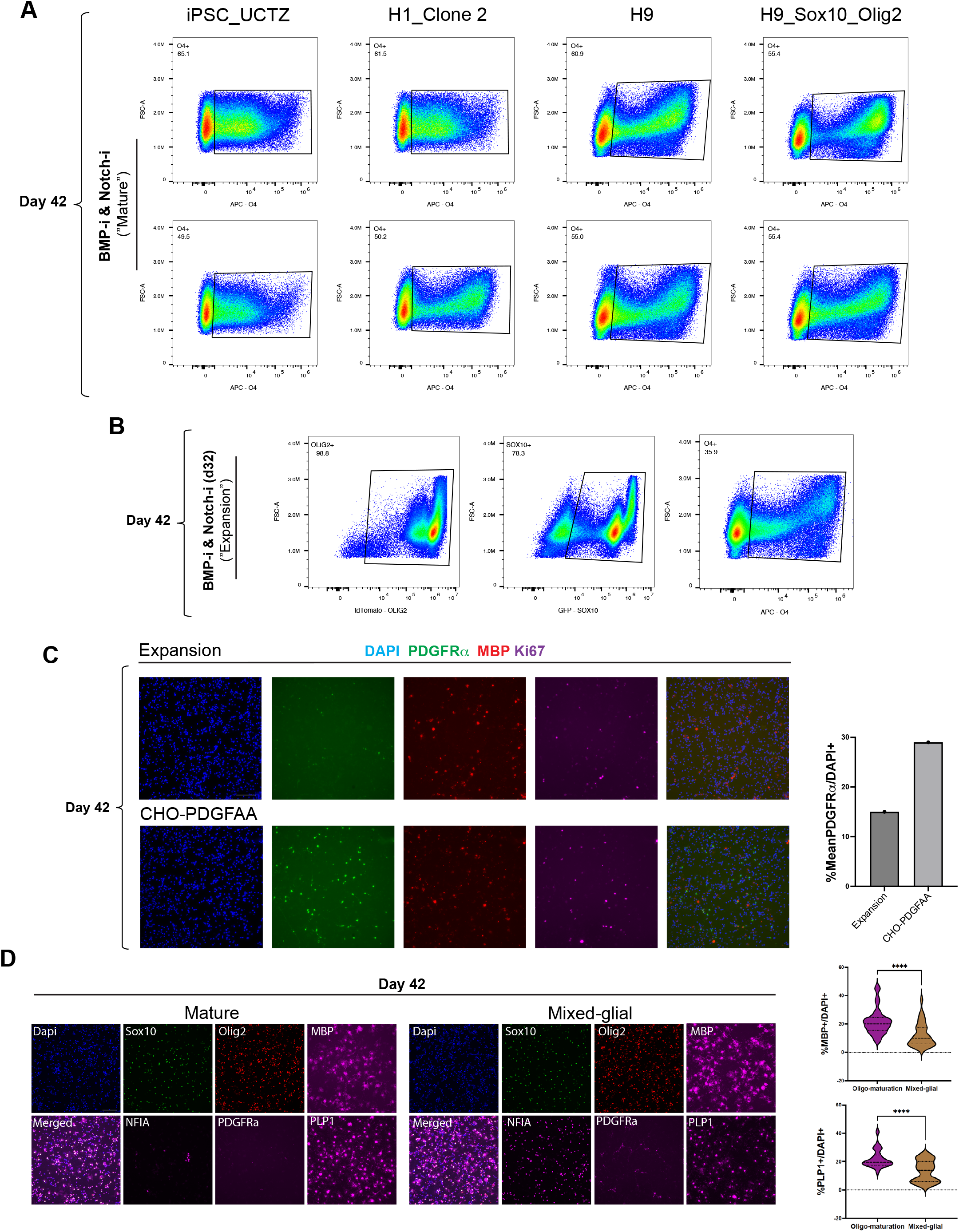
The dual inhibition of NOTCH and BMP allows the maturation of oligodendrocyte across multiple human pluripotent stem cell lines, while early NOTCHi enables the expansion of a PDGFRα cycling population. (A) FACS plots show the O4+ percentage across iPSC and ESC cell lines using the mature protocol, which involves the dual inhibition of BMP and NOTCH from day 26 to day 42. (B) Delayed NOTCHi in the expansion protocol enables the expansion of the SOX10+ population while slowly converting them to more mature O4+ expressing oligodendrocytes. (C) Representative immunofluorescence images show an expansion of a PDGFRα+ cycling population in organoids treated with early NOTCHi from day 26 to day 32, and then with DMH1, FGF18 and a more potent human recombinant PDGF-AA protein at 20ng/mL (R&D #11564-PA) (CHO vs Expansion conditions). (D) Quantification of MBP+ and PLP1+ cells in mature and mixed-glial protocol. Scale bar: 100μm.

**Supplementary Fig 5.**
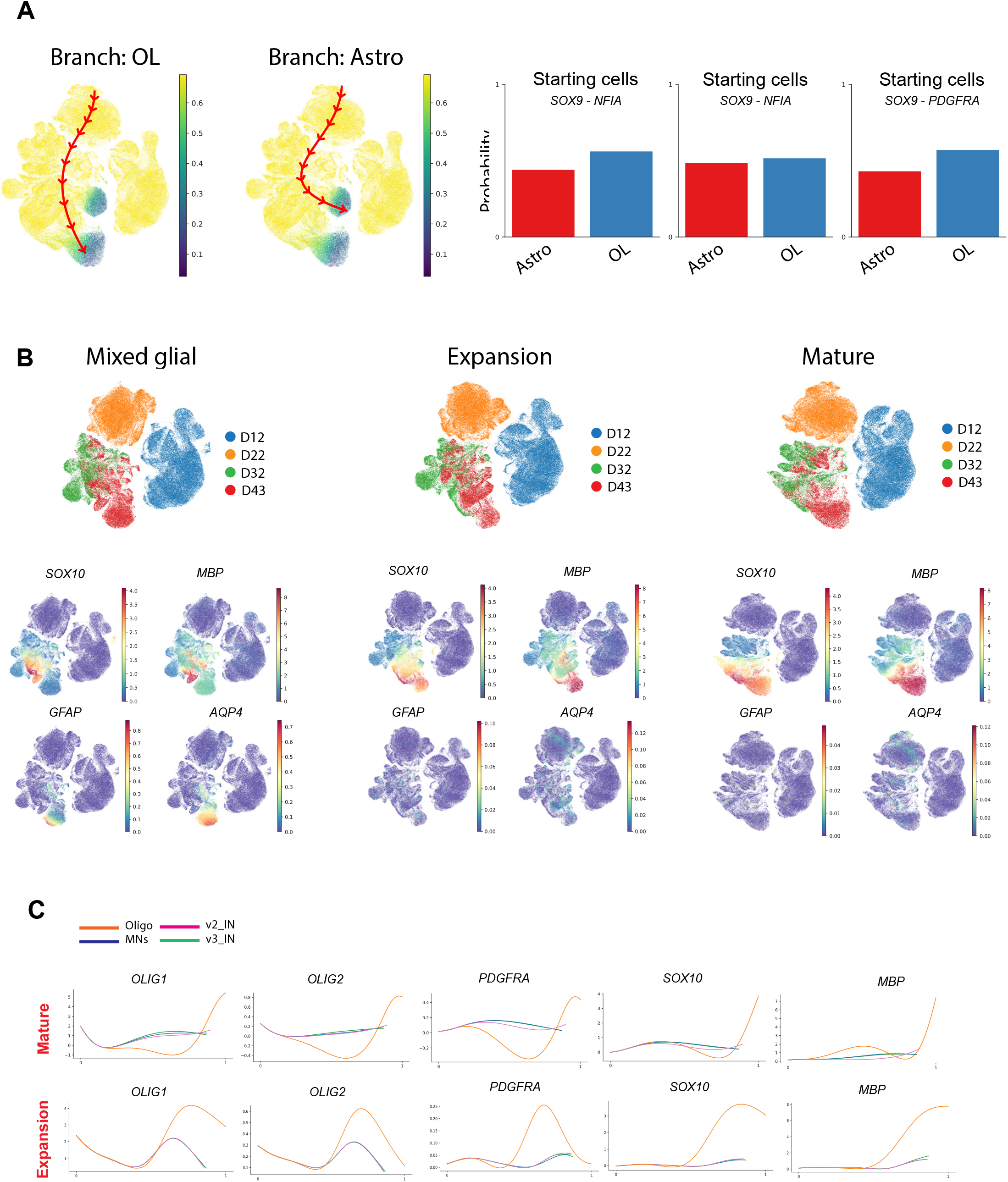
Lineage trajectory analysis reveals time-dependent differences in glial development across three protocols. (A) Trajectory reconstruction using Palantir of both oligodendrocytes and astrocytes across our four time-points (left). The probability bar plots show how likely a day 22 and 32 cell positive for SOX9-NFIA, and a day 32 NFIA-PDGFRα become either an oligodendrocyte or an astrocyte (right). (B) Palantir-based clustering show canonical oligodendrocyte and astrocyte markers expression in mature, expansion, and mixed-glial protocols. (C) The delayed NOTCHi from day 26 to day 32 enables an overall delay of developmental steps across oligo differentiation.

**Supplementary Fig 6.**
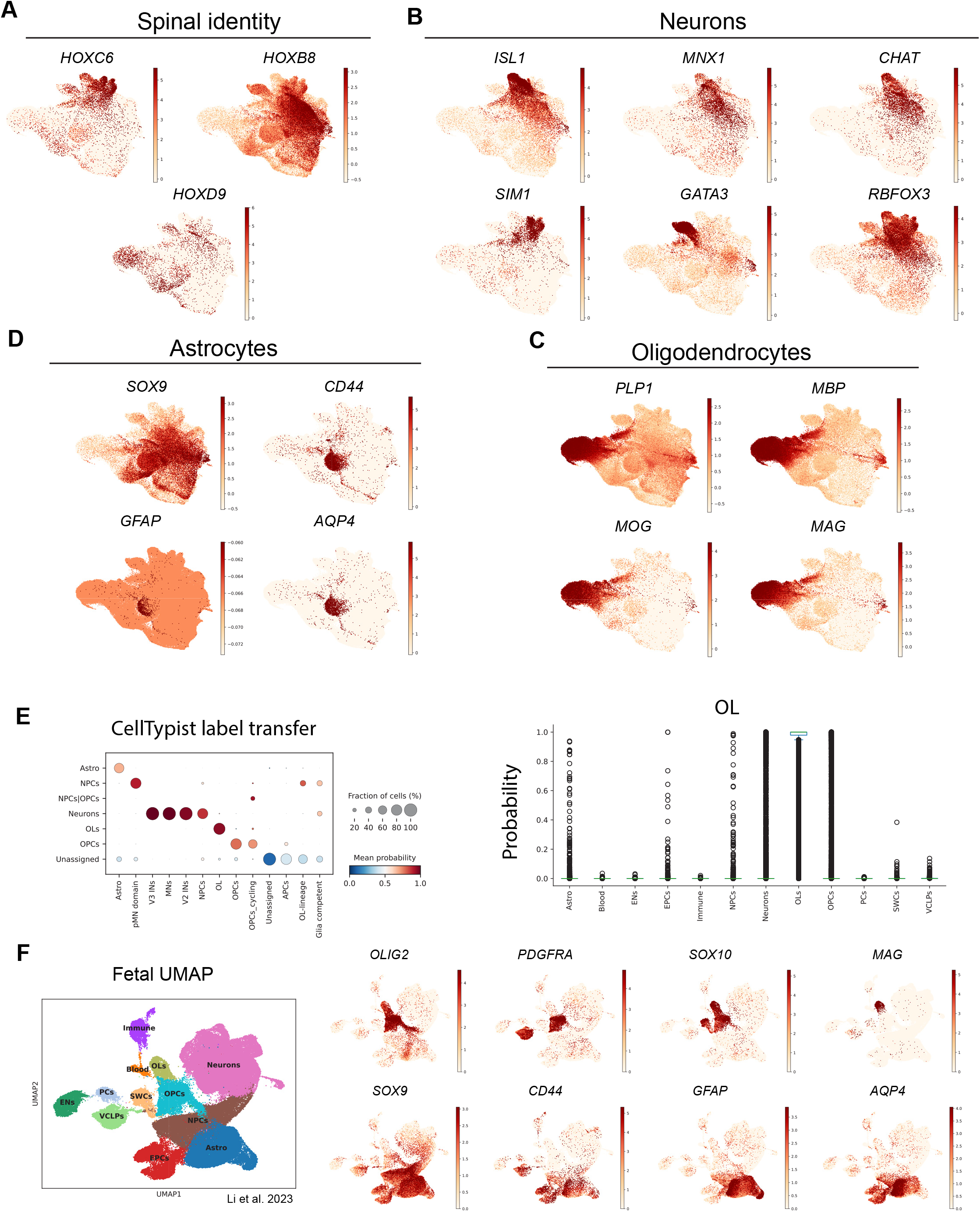
Single cell RNA-sequencing data identifies oligodendrocytes, astrocytes and neurons across the three protocols closely matching corresponding cell types in human fetal data. Gene expression plots showing spinal cord identity (A), neuron (B), oligodendrocyte (C), and astrocyte markers (D). (E) Characterization of OPCs, oligodendrocytes and astrocytes within the human fetal spinal cord dataset published by Li et al^48^, with re-annotation of the *CD44-GFAP-APQ4* cluster as “Astro” (astrocytes). (F) Dot plot (left) showing CellTypist label transfer, highlighting the high probability and the relative fraction of cells from our *in vitro* data closely match the cell types in the published human fetal spinal cord atlas. Box plot (right) showing CellTypist oligodendrocyte prediction scores across annotated cell types, identifying very high probability match.

**Supplementary Fig 7.**
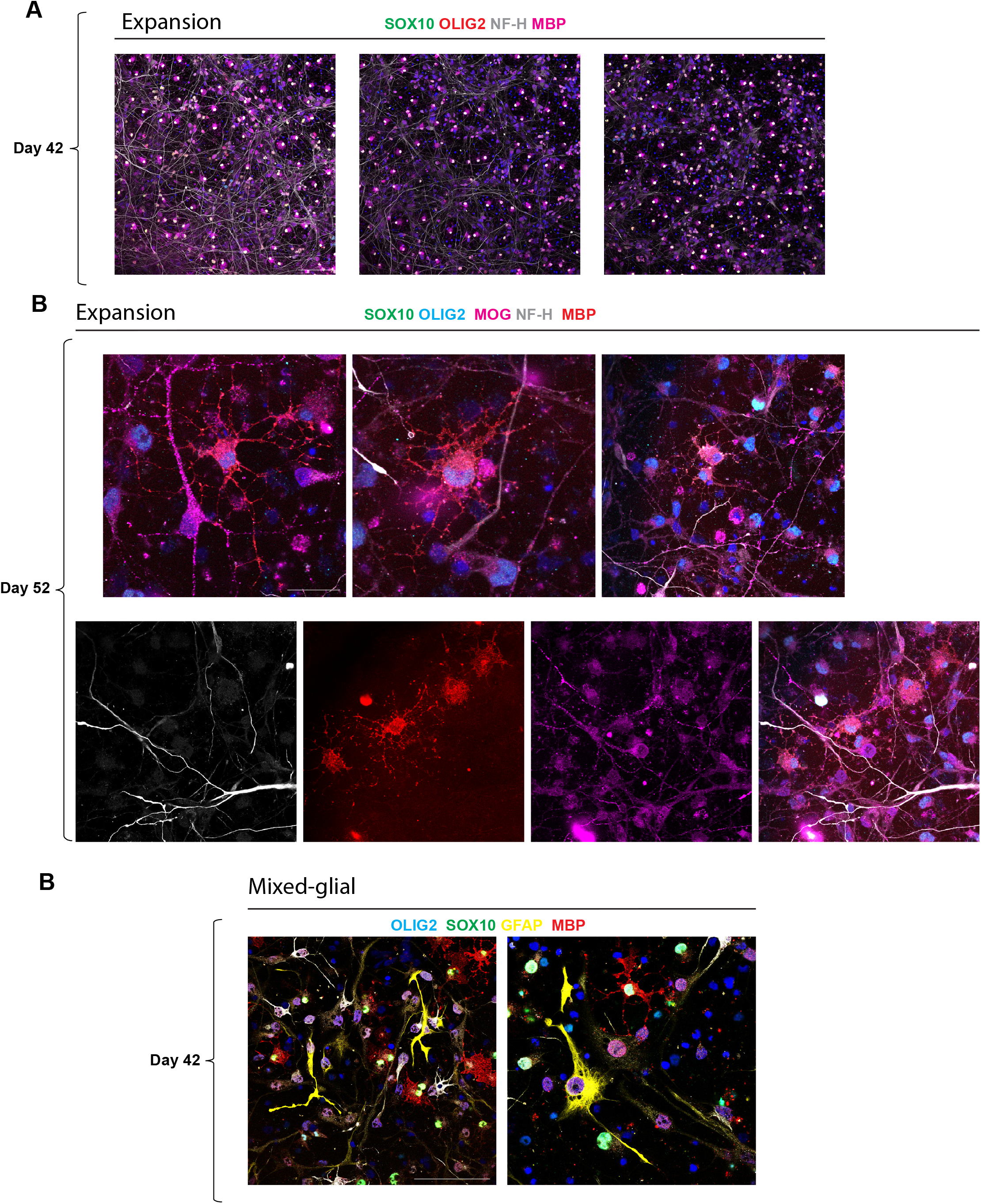
Identification of mature oligodendrocytes in the *expansion* and *mixed-glial* protocols. Representative immunofluorescence images of the *expansion* protocol show day 42 (A) and day 52 (B) oligodendrocytes positive for MOG and MBP that closely align with NF-H neuronal axons. (C) Characterization of both oligodendrocytes positive for MBP and astrocytes positive for GFAP in the *mixed-glial* protocol. Scale bar: 100μm.

